# Single-cell integration and multi-modal profiling reveals phenotypes and spatial organization of neutrophils in colorectal cancer

**DOI:** 10.1101/2024.08.26.609563

**Authors:** Valentin Marteau, Niloofar Nemati, Kristina Handler, Deeksha Raju, Alexander Kirchmair, Dietmar Rieder, Erika Kvalem Soto, Georgios Fotakis, Sandro Carollo, Nina Boeck, Alessia Rossi, Alexandra Scheiber, Arno Amann, Andreas Seeber, Elisabeth Gasser, Steffen Ormanns, Michael Günther, Agnieszka Martowicz, Zuzana Loncova, Giorgia Lamberti, Anne Krogsdam, Michela Carlet, Lena Horvath, Marie Theres Eling, Hassan Fazilaty, Tomas Valenta, Gregor Sturm, Sieghart Sopper, Andreas Pircher, Patrizia Stoitzner, Peter J. Wild, Patrick Welker, Pascal J. May, Paul Ziegler, Markus Tschurtschenthaler, Daniel Neureiter, Florian Huemer, Richard Greil, Lukas Weiss, Marieke Ijsselsteijn, Noel F.C.C. de Miranda, Dominik Wolf, Isabelle C. Arnold, Stefan Salcher, Zlatko Trajanoski

**Affiliations:** Biocenter, Institute of Bioinformatics, Medical University of Innsbruck, Austria; Institute of Experimental Immunology, University of Zurich, Switzerland; Department of Internal Medicine V, Haematology & Oncology, Comprehensive Cancer Center Innsbruck and Tyrolean Cancer Research Institute, Medical University of Innsbruck, Austria; Department of Visceral, Transplant and Thoracic Surgery, Medical University Innsbruck, Austria; Innpath, Tirol Kliniken, Medical University Innsbruck, Austria; Tyrolpath Obrist Brunhuber GmbH, Zams, Austria; Department of Therapeutic Radiology and Oncology, Medical University Innsbruck, Austria; Department of Molecular Life Sciences, University of Zürich, Switzerland; Institute of Molecular Genetics of the Czech Academy of Sciences, Prague, Czech Republic; Department of Dermatology, Venereology and Allergology, Medical University of Innsbruck, Austria; Senckenberg Institute of Pathology, Goethe University Frankfurt, Germany; University Cancer Center Frankfurt (UCT), Germany; Translational Cancer Research and Institute of Experimental Cancer Therapy, Klinikum rechts der Isar, School of Medicine & Health, Technical University of Munich, Germany; Institute of Pathology, Paracelsus Medical University, Salzburg, Austria; Department of Internal Medicine III, Paracelsus Medical University, Salzburg, Austria; Austrian Breast & Colorectal Cancer Study Group (ABCSG), Vienna, Austria; Department of Pathology, Leiden University Medical Center, The Netherlands; Comprehensive Cancer Center Zürich, Zürich, Switzerland; Institute of Pathology, Neuropathology and Molecular Pathology, Medical University of Innsbruck, Austria; Boehringer Ingelheim International Pharma GmbH & Co KG, Biberach, Germany

**Keywords:** Colorectal cancer, single-cell sequencing, spatial single-cell profiling, single-cell atlas, cellular niches

## Abstract

The immune composition of the tumor microenvironment (TME) has a major impact on the therapeutic response and clinical outcome in patients with colorectal cancer (CRC). Here, we comprehensively characterize the TME at the single-cell level by first building a large-scale atlas that integrates 4.27 million single cells from 1,670 patient samples. We then complemented the atlas with single-cell profiles from four CRC cohorts with 266 patients, including cells with low mRNA content, spatial transcriptional profiles from 3.7 million cells, and protein profiles from 0.7 million cells. The analysis of the atlas allows refined tumor classification into four immune phenotypes: immune desert, B cell enriched, T cell enriched, and myeloid cell enriched subtypes. Within the myeloid compartment we uncover distinct subpopulations of neutrophils that acquire new functional properties in blood and in the TME, including anti-tumorigenic capabilities. Further, spatial multimodal single-cell profiling reveals that neutrophils are organized in clusters within distinct functional niches. Finally, using an orthotopic mouse model we show that cancer-derived systemic signals modify neutrophil production in the bone marrow, providing evidence for tumor-induced granulopoiesis. Our study provides a big data resource for the CRC and suggests novel therapeutic strategies targeting neutrophils.

## INTRODUCTION

Colorectal cancer (CRC) is the most prevalent gastrointestinal malignancy and represents the second leading cause of cancer-related mortality worldwide according to the GLOBOCAN data^1^. Despite the improvement of early detection screening policies and the development of many novel treatments, the median overall survival (OS) for patients with metastatic CRC (mCRC) is ∼30 months^2^. Advancements in targeted therapy for mCRC have been limited, especially when contrasted with the significant progress achieved in other solid tumors such as non-small cell lung cancer^3^. The genetic heterogeneity^4^ as well as paucity of drugs targeting common driver genes, pose considerable challenges for developing precision oncology approaches for patients with CRC. Moreover, CRC appears to exhibit resistance to immune checkpoint inhibitor (ICI) therapy, with the notable exception of approximately 4% of metastatic cases characterized by a damaged mismatch repair system (dMMR) or/and high microsatellite instability (MSI-H)^5,6^. This is somehow paradoxical since CRCs are known to be under immunological control, as we have shown in the past^7^. Thus, there is an urgent need to explore new immunomodulatory strategies for the vast majority of CRC patients with microsatellite stable (MSS) tumors. Strategies that convert the “immune cold” tumor microenvironment (TME) to “immune hot” could sensitize cancers to ICI therapy. A number of clinical trials with drug combinations have been carried out to overcome resistance to immunotherapy in MSS CRC^8,9^. The benefits of using a dual combination of ICIs (CTLA-4 and PD-1 blockade) or adding ICIs to chemotherapy appear to be very limited^8,10^. Combination strategies that exploit synergistic potential of ICI and targeted drugs showed increase of the anti-tumor activity, however, the objective response rates were disappointingly low^8^. Thus, to address this unmet clinical need a better understanding of the immunosuppressive TME and its heterogeneity is of utmost importance.

The intrinsic complexity of the interaction of the two interwoven systems, the tumor and the immune system, poses considerable challenges and requires comprehensive approaches to interrogate cancer immunity during tumor initiation and progression, and following therapeutic modulation thereof. A number of established and novel high-throughput technologies enable the generation of the necessary data and thereby provide the basis for mechanistic understanding. Specifically, next-generation sequencing (NGS) of cancer samples provide large data sets that can be mined for immunologically relevant parameters (see our reviews^11,12^). Using these tools we were able to chart the TME in CRC^13^ and developed The Cancer Immunome Atlas^14^. While these attempts uncovered the enormous heterogeneity of MSS CRC and revealed unique cancer cell and TME subtypes with prognostic and therapeutic relevance, the advent of single-cell RNA sequencing (scRNA-seq) technologies opened fundamental new avenues to dissect cancer heterogeneity. We recently developed a novel computational approach to dissect the cellular diversity in the TME at high resolution by integrating expression profiles of single cells from multiple studies using cutting-edge artificial intelligence (AI) methods^15^. Similar effort was recently carried out in the context of CRC^16^, enabling stratification of patients. However, limited by the sampling bias of the commonly used 10x Genomics platform, this atlas is lacking neutrophils. Neutrophils are the most abundant cells circulating in the human body and are emerging as important regulators of cancer^17–20^. Neutrophils are highly abundant in intratumoral as well as stromal regions in CRC^18^ and co-localize with CD8^+^ T cells^19^. Neutrophils exhibit remarkable plasticity, show different pro-tumor and anti-tumor functions^20^, and can acquire anti-tumor phenotype in response to immunotherapy^21^. This dual role of the neutrophils in the TME is a consequence of the intracellular interactions with cancer cells as well as with other components of the TME, including T cells, macrophages and endothelial cells^22^. The importance of neutrophil-T cell interactions is supported by the observation that CD4^+^ PD-1^+^ T cells in granulocyte-dominated cell neighborhoods are associated with favorable prognosis in CRC^23^. Hence, deep characterization of neutrophils in the TME, in normal tissue, and blood from CRC patients would provide novel insights into the myeloid compartment and the myeloid-lymphoid cell-cell interactions.

Here, we first integrated 76 CRC single-cell RNA sequencing datasets spanning 4.27 million cells from 650 patients and 1,670 samples. We then complemented the atlas by profiling single cells from four cohorts (n=266 patients) including: 1) single-cell profiling of samples using an optimized protocol and platform that efficiently recovers low mRNA-content cells; 2) spatial single-cell transcriptional profiling of 3.7 million cells as well as spatial single-cell protein profiling of 715,000 cells. The analyses refined tumor classification into four immune phenotypes. Within the myeloid compartment we uncovered distinct subpopulations of neutrophils that acquire new functional properties in the TME including antigen-presenting capabilities (HLA-DR^+^ neutrophils). Furthermore, spatial multimodal single-cell profiling revealed that neutrophils are organized in clusters within distinct multi-cellular niches. Finally, using an orthotopic mouse model we show that cancer-derived systemic signals modify neutrophil production in the bone marrow, providing evidence for tumor-induced granulopoiesis.

## RESULTS

### Generation of a large-scale CRC single-cell atlas

Using our approach for the integration of single-cell datasets^15^ we first compiled CRC related scRNA-seq data from 48 studies and 76 datasets comprising 4.27 million cells from 650 patients and 1,670 samples (Figure 1A). This comprehensive CRC single-cell atlas integrates curated and quality-assured transcriptomic data from publicly available studies, complemented with extensive metadata curation, and encompasses the full spectrum of disease progression, from normal colon to polyps, primary tumors, and metastases, covering both early and advanced stages of CRC (see STAR Methods, Figures 1B and C and Figure S1A-E). Study characteristics are summarized in Suppl. Table S1. In total, the atlas includes transcriptomic data from 487 CRC patients (median: 58 years [22-91]), 99 patients with polyps (median: 60 years [40-75]), and 64 control individuals (median: 57 years [23-79]). Of the 487 CRC patients, 254 had colon cancer (COAD), 63 rectal cancer (READ), and 170 were not otherwise specified. Of these, 89 were MSI-H, 268 MSS, and 130 had an unknown microsatellite status. In terms of clinicopathological disease stage, 41 patients were classified as TNM stage I, 121 as stage II, 171 as stage III, and 110 as stage IV, while the stage was not reported for 44 patients. CRC samples include tissue of the primary tumor (n=674) or CRC liver metastasis (n=175) that were obtained by surgical resection. Among the control samples, 284 were derived from distant non-malignant tissue of colon tumor patients, of which 220 have a patient-matched tumor sample. 186 non-tumor control samples as well as 116 polyps and 90 matched normal samples were obtained from patients undergoing colonoscopy. Further, 103 blood samples of CRC patients as well as 24 samples derived from tumor affected lymph nodes, with another 18 non-tumor affected lymph node samples were included. Of the control patients none had a history of inflammatory bowel diseases.

**Figure 1.**
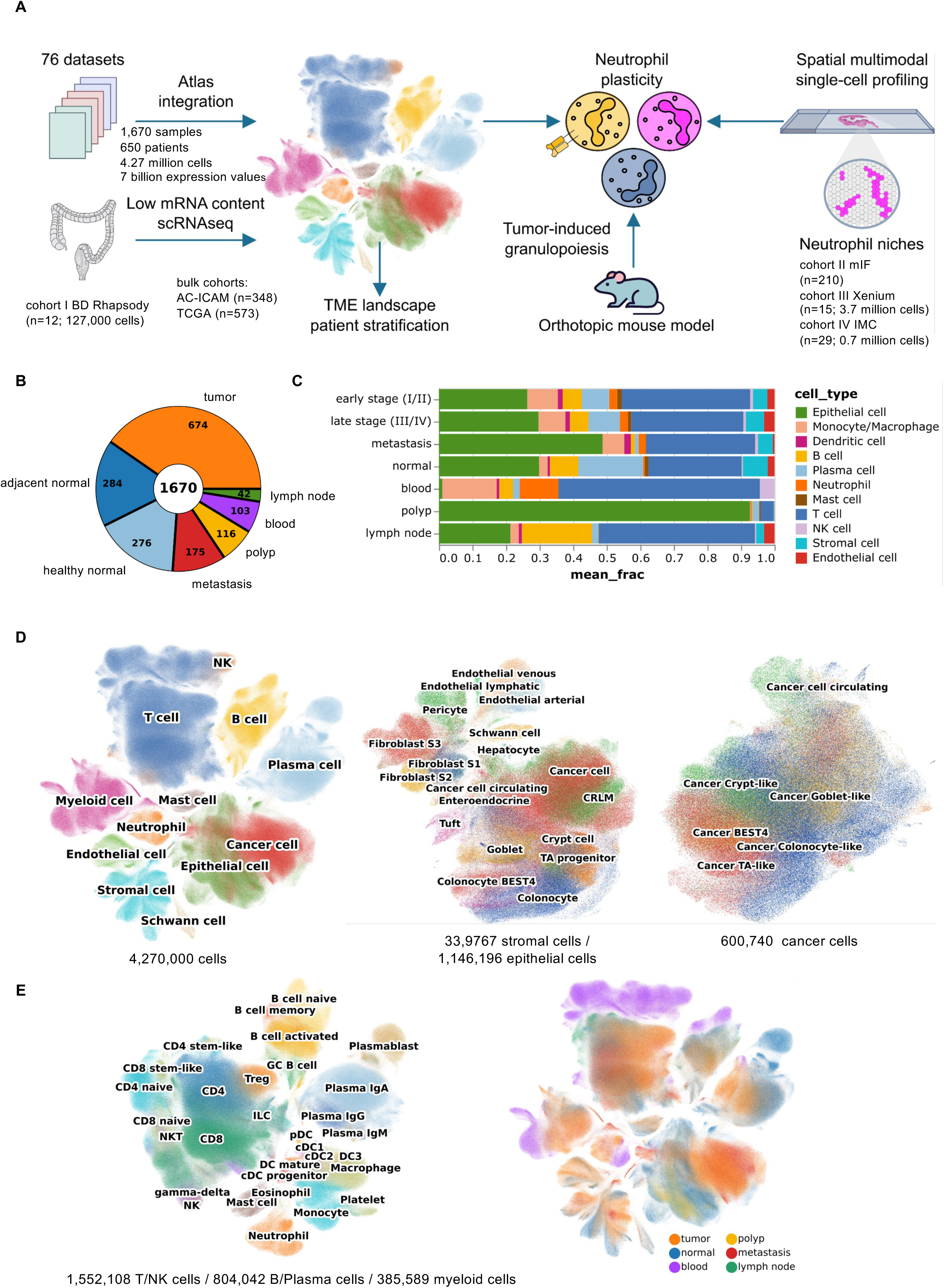
Schematic outline of the overall concept of the study including publicly available and generated datasets. (**A**) Summary of the data integration and analysis workflow. (**B**) Number of samples for each type. (**C**) Overview of the cell fractions in different tissues. (**D**) Overview of the CRC atlas and the epithelial, and stromal/endothelial components depicted as UMAP plots. (**E**) Overview of the CRC atlas sample types and the immune components depicted as UMAP plots.

Although neutrophils play a prominent role in cancer biology, they are underrepresented in most available scRNA-seq datasets generated with the widely used 10x Genomics technology^24^ (Figure S1 F). To address this, we generated scRNA-seq data from peripheral blood, adjacent normal tissue, and tumor tissue from CRC patients from cohort I (n = 12) using the BD Rhapsody platform and our optimized protocol^25^ that allows capture of cells with very low mRNA content^24^. The data was then integrated into the final high-resolution single-cell atlas for CRC. Overall, the atlas integrates 4,264,946 single cells, which are annotated to 12 coarse cell type identities and 62 major cell subtypes or cell states (e.g. dividing cells) based on previously established canonical single-cell signatures including 1,146,196 epithelial cells, 1,552,108 T/NK cells, 804,042 B/Plasma cells, 385,589 myeloid cells, and 33,9767 stromal and endothelial cells (Figure 1D). Cancer cells (in total 600,740 cells, from primary tumor and metastatic tissue) in the atlas showed high heterogeneity in their transcriptomic profiles (Figure 1D). Due to the large patient cohort, we were able to apply high-resolution cancer cell classification based on their specific marker gene expression signatures (Figure S1D). We divided the following main clusters: 1) colonocyte-like (*FABP1, CEACAM7, ANPEP, SI*), 2) BEST4 (BEST4*, OTOP2, SPIB*), Goblet-like (*MUC2, FCGBP, ZG16, ATOH1*), 3) Crypt-like (*LGR5, SMOC2*), and 4) transit-amplifying-like (*TOP2A, UBE2C, PCLAF, HELLS, TK1*) (Figure 1D, Figure S1C). There are highly mitotic/proliferative clusters (*LGR5, TOP2A, MKI67*) of both COAD and READ that resemble highly aggressive and invasive cancer cells. In addition, *EPCAM* expressing cells in the blood of CRC patients were annotated as circulating cancer cells.

With a total of 2,741,739 immune cells, this single-cell atlas enables high-resolution classification of immune cell types across both innate and adaptive subsets (Figure 1C, Figure S1A-C). These include T cells -CD4 (*CD4*), CD8 (*CD8A*), T regulatory cells (*FOXP3*), and NK cells (*NKG7*, *TYROBP*). B cells are represented as naïve (*IGHD*, *TCL1A*), memory (*CD27*,*CD44*), and activated (*CD69*, *CD83*) subsets, while plasma cells (*JCHAIN*) express immunoglobulins such as IgA (*IGHA1*), IgG (*IGHG1*), and IgM (*IGHM*). The myeloid lineage encompasses monocytes (*VCAN*, *CD14*), macrophages (*C1QA*, *C1QB*), mast cells (*TPSB2*, *CPA3*), dendritic cells, including conventional DC type 1 (*cDC1*, *CLEC9A*, *XCR1*) and cDC2 (*CD1C*, *CLEC10A*), and plasmacytoid DC (*pDC*, *IL3RA*, *GZMB*), as well as eosinophils (*CLC*, *CCR3*) and neutrophils (*CXCR2*, *FCGR3B*). Specialized populations such as gamma-delta T cells (*TRDV1*, *TRGC1*) and innate lymphoid cells (ILCs; *IL4I1*, *KIT*) are also depicted, along with immune cell developmental stages, such as progenitors (*CD24*, *AXL*) and stem-like cells (*LEF1*), and cycling states (*MKI67*, *CDK1*).

### TME characterization at the single-cell level enables stratification of tumors into distinct immune phenotypes

Utilizing the high-resolution atlas we stratified 284 CRC patients with primary tumor samples based on the infiltration patterns of their TME. The high number of patients enabled us to perform unsupervised clustering of batch-corrected cell-type fractions, revealing four major immune clusters with distinct subclusters (Figure 2A and Figure S2A): 1) immune deserted tumors (ID; i.e. no significant immune cell infiltration but high cancer cell fraction); 2) the subtype of tumors with T cell dominance (T; CD8^+^, CD4^+^, T regulatory cells); 3) the subtype of tumors with B cell dominance (B; B cell, plasma IgA and IgG cell) and 4) the subtype of tumors with myeloid dominance (M; macrophage/monocyte). Using logistic regression, we did not find any association of the four different patient strata to tumor stages (early I/II *vs.* advanced III/IV), sex, or other features. However, patients in the M (p=0.013) or B (p=0.089) cluster were associated with a higher likelihood of being in an early tumor stage (I/II). In addition, the T cell cluster (p=0.026) subcluster was associated with a higher likelihood of the tumor being MSI-H compared to MSS.

**Figure 2.**
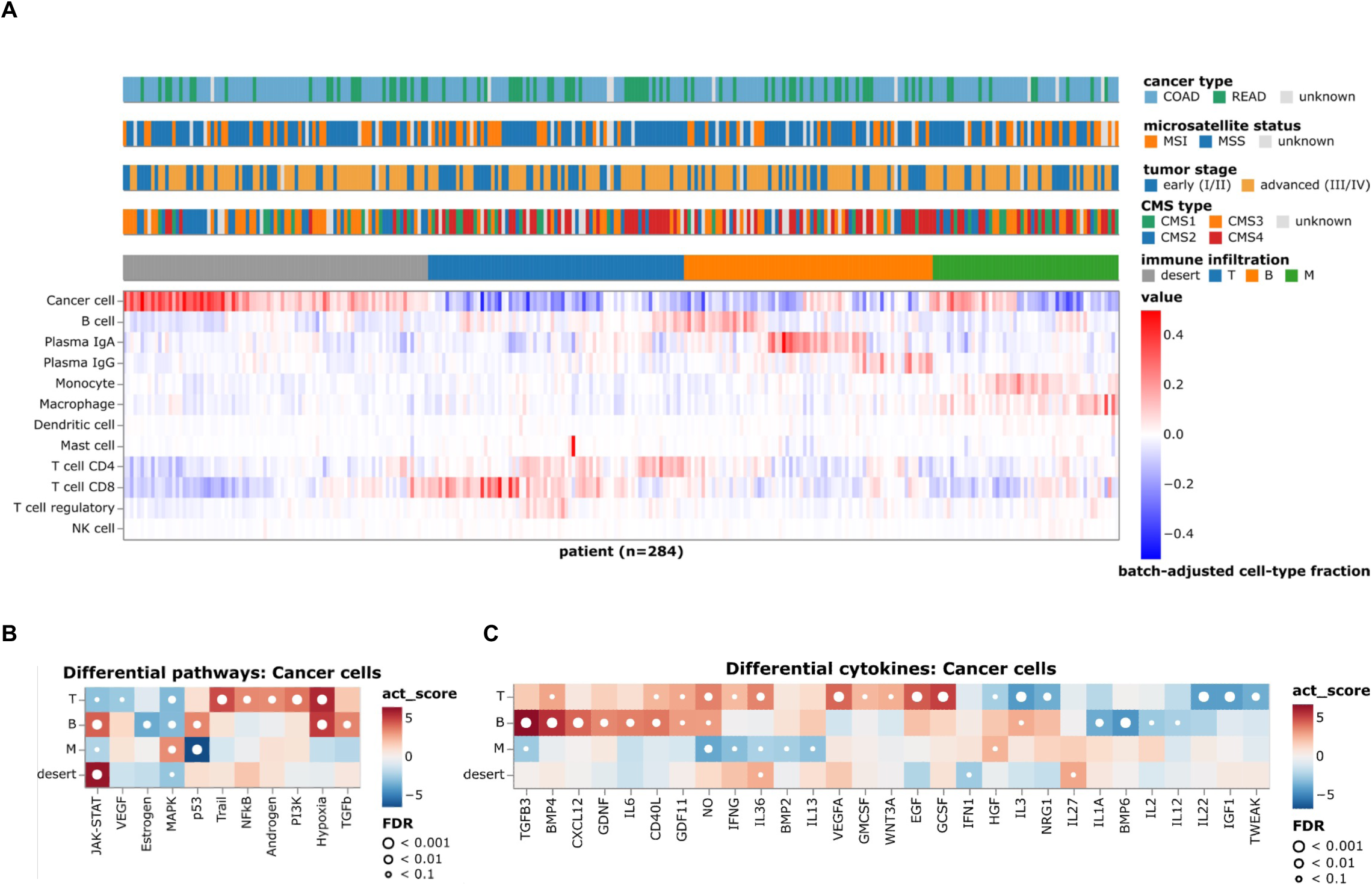
CRC patient stratification based on tumor immune phenotypes. (**A**) Patient characteristics and stratification of the tumor immune phenotypes. (**B**) Differential activation of PROGENy cancer pathways across different cell types in the TME. Heatmap colors indicate the deviation from the overall mean, independent of tumor histology and stage. White dots indicate significant interactions at different false-discovery-rate (FDR) thresholds. Only cytokine signatures with an FDR < 0.1 in at least one patient group are shown. (**C**) CytoSig cytokine signaling signatures in cancer cells between the four tumor immune phenotypes. Heatmap colors indicate the deviation from the overall mean, independent of tumor histology and stage. White dots indicate significant interactions at different FDR thresholds. Only cytokine signatures with an FDR <0.1 in at least one patient group are shown

In order to identify tumor-cell based TME imprinting characteristics, we next analyzed differentially enriched pathways in the cancer cells of each immune infiltration type using the tool PROGENy^26^ (Figure 2B and S2B). The ID subtype displayed an upregulation (false discovery rate [FDR] < 0.1) of the JAK-STAT pathway (Figure 2B). The p53 pathway was significantly downregulated (false discovery rate [FDR] <0.1) in the M subtype. We then used the CytoSig tool^27^ to define enriched cytokine signaling signatures in cancer cells of each immune subtype (Figure 2C and S2C). CytoSig platform uses transcriptomics data to define cytokine signatures and signaling activities in order to investigate mechanisms behind diseases where cytokines are involved. The signature of the transforming growth factor beta-3 (TGFB3) cytokine was significantly elevated in the B subtype (false discovery rate [FDR] <0.1) (Figure 2C and S2C). The majority of the signatures were downregulated in the ID subtype (Figure 2C and S2C).

### Integration of bulk RNA-seq data reveals genotype-immune phenotype associations

The majority of the scRNA-seq studies lack both comprehensive cancer genotype information and survival data. In contrast, the TCGA^4^ or the more recent AC-ICAM^28^ include this information, alongside bulk RNA-sequencing data. Building on our previous work that identified genomic features influencing immune phenotypes in a pan-cancer study using bulk RNA-seq data^14^, we leveraged the high resolution of the single-cell CRC atlas for a more detailed analysis of these determinants. We first developed Shears, a novel computational method to address major limitations of existing algorithms for associating single cells from scRNA-seq datasets with bulk RNA-seq phenotypic or clinical information, such as computational inefficiency on large datasets, and the inability to account for covariates and biological replicates (see Methods). We then combined bulk RNA-seq data from the TCGA (n=573) and the AC-ICAM (n=348) cohorts and used Shears to assess the relationships between survival outcomes as well as specific genotypes with atlas-derived cell-type transcriptomic signatures (Figure 3A). Overall, the presence of eosinophils, plasma cells and B cells, as well as several T cell subtypes including CD8^+^ T cells, CD4^+^ T cells, and γδ T cells was associated with improved survival in CRC patients. Conversely, CRC patients with tumors enriched with endothelial cells, stromal cells, and neutrophils were associated with worse survival outcomes (Figure 3A). However, distinct tumor genotypes like *BRAF* and *KRAS* mutated tumors were associated with different immune infiltration patterns (Figure 3B-C). For example, tumors from CRC patients with *KRAS* mutations had increased infiltrations of neutrophils and eosinophils whereas the same cell types were depleted in tumors from CRC patients with *BRAF* mutations. Similarly, plasma cells were depleted in tumors with *APC* mutations (Figure S3A). Conversely, *TP53* mutations were linked with infiltration of plasma cells (Figure S3B). This single-cell perspective of the TME strengthens the link between the tumor’s genetic profile, its histological features, and the associated immune contexture^29^. Given the importance of the driver genes in terms of treatment decisions, this finding could be exploited to derive rationale for personalized therapeutic combination strategies based on the underlying genetic tumor profile.

**Figure 3.**
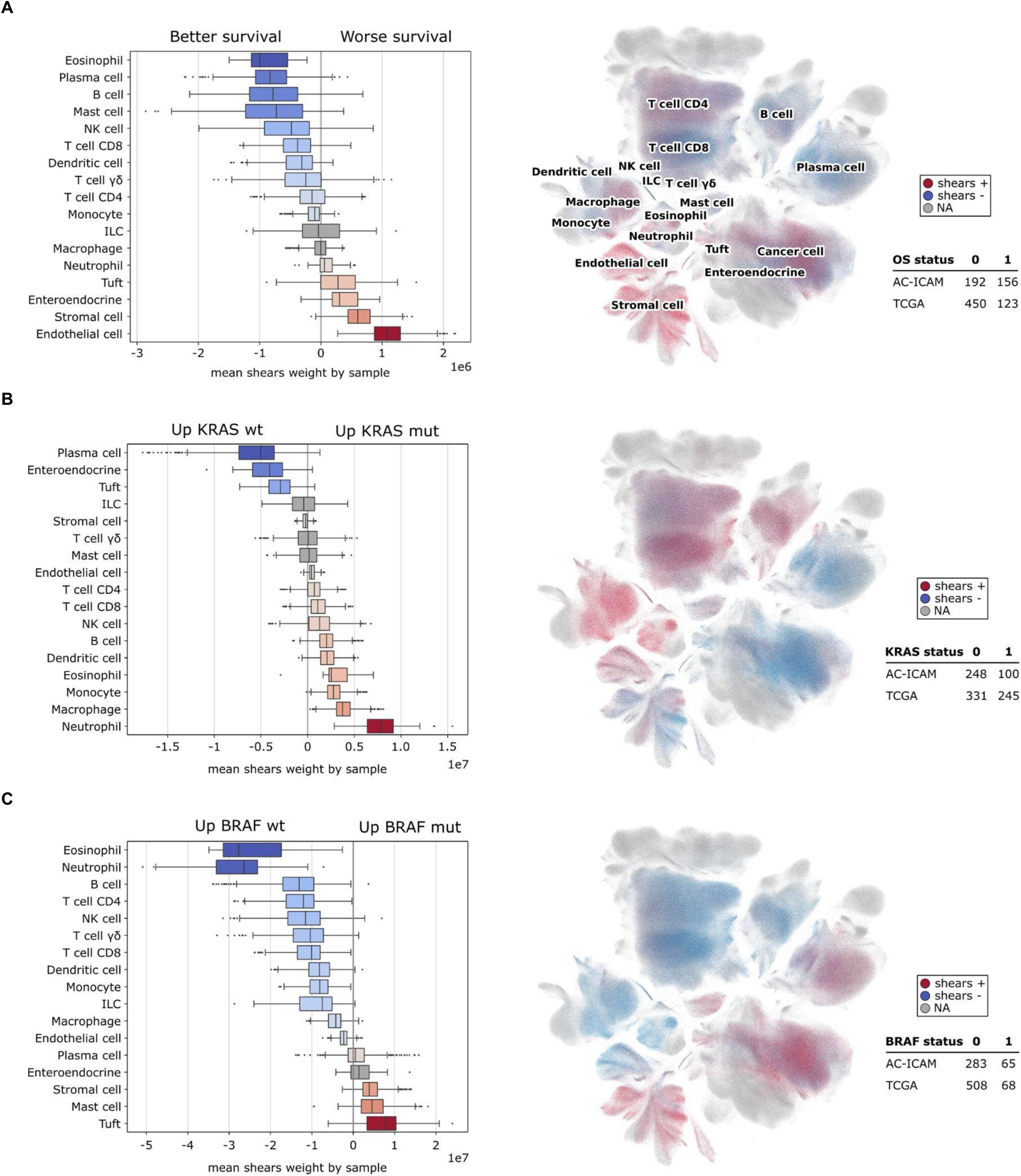
Associations of cellular composition, survival, and genotypes. (**A-C**) *SHEARS* analysis showing the association of cellular composition and phenotypic information from bulk RNA-seq data from the combined AC-ICAM and TCGA cohorts. Boxplots summarize the central Shears tendencies for each cell type in relation to the phenotype, with each data point representing the mean Shears weight/coefficient by patient and cell type with min 10 cells. UMAP plots indicate the position of cells positively (red) or negatively (blue) associated with mutation or better survival. (**A**) Association of cellular composition with overall survival. (**B**) Association of cellular composition with KRAS mutation in CRC patients (n = 345). (**C**) Association of cellular composition with BRAF mutation in CRC patients (n = 133).

### Circulating and tumor-associated neutrophils have diverse and plastic phenotypes

Given the lack of single-cell data for neutrophils, we carried out comprehensive characterization of the data we generated using the BD Rhapsody platform. The obtained scRNA-seq dataset comprises 126,991 single cells categorized into 23 main cell types based on specific marker genes, including 55,245 epithelial/tumor cells, 4,428 stromal/endothelial cells, and 65,782 immune cells, including 30,924 neutrophils (Figure 4A). Compositional analysis using scRNA-seq and flow cytometry showed high concordance when comparing blood, normal, and tumor tissue samples (Figure 4B). As expected, neutrophils were the most abundant leukocyte subtype in blood samples (60–70%), followed by T lymphocytes and monocytes (Figure 4B). Normal-adjacent tissue was dominated by epithelial cells, while tumor tissue contained a higher proportion of immune cells. Notably, neutrophils comprised ∼20% of all cells and ∼40% of leukocytes in tumor tissue but were present only in small numbers in normal-adjacent tissue, which was confirmed using flow cytometry (Figure 4C). Within the granulocytes lineage, neutrophils were the most abundant cells in the tumor and in blood (95.2% and 96.1%, respectively), whereas in normal adjacent tissue the fractions for neutrophils (36.8%), eosinophils (35.6%) and mast cells (27.6%) were similar.

**Figure 4.**
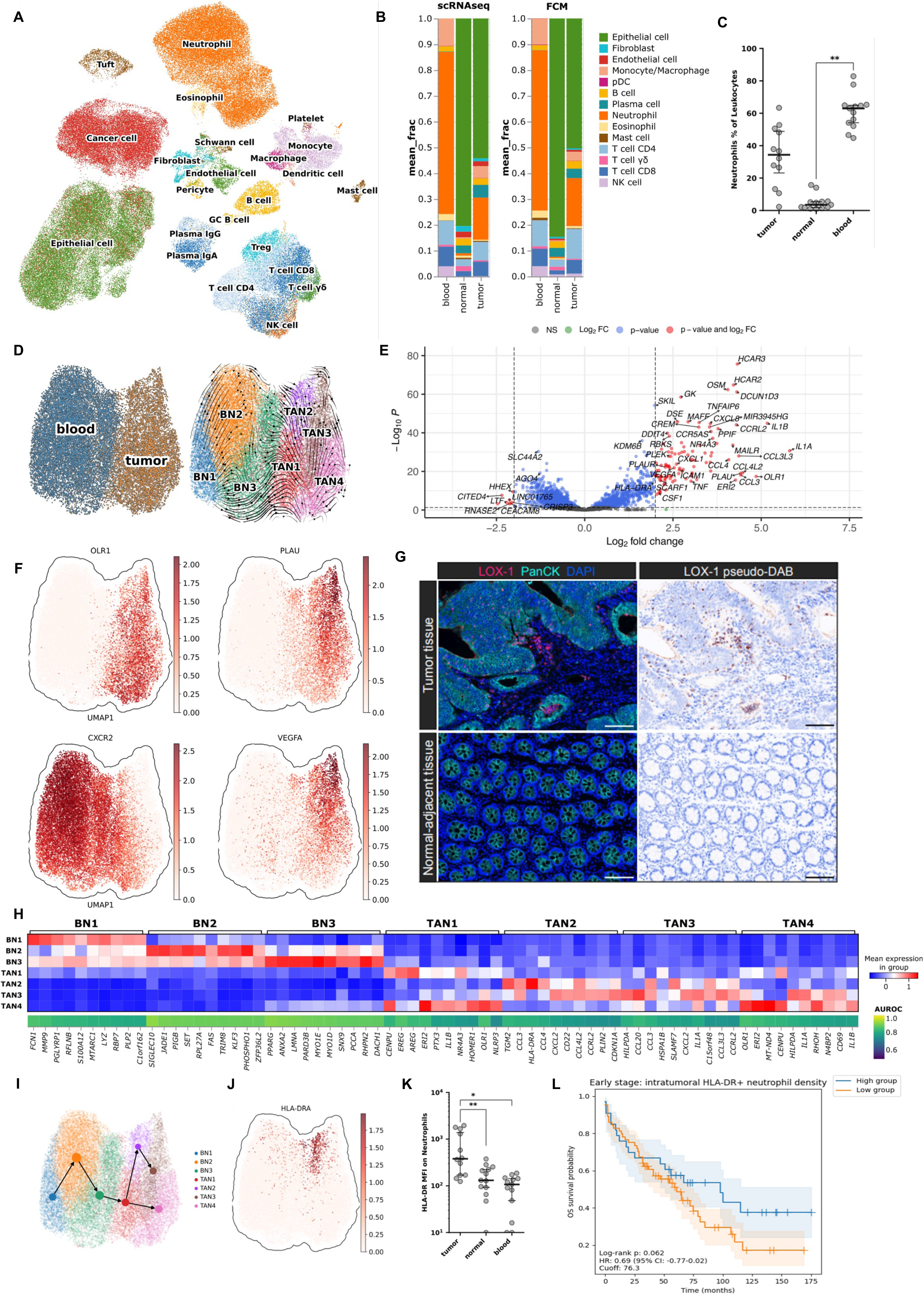
Characterization of neutrophils in CRC. (**A**) UMAP of the generated CRC scRNA-seq dataset (cohort I; n = 12 patients; tumor tissue, normal-adjacent tissue and matched blood) colored by cell type. (**B**) Cell-type fractions obtained by scRNA-seq and by flow cytometry. (**C**) Neutrophil fractions (as percentage of leukocytes) obtained by flow cytometry (n = 12 patients; tumor tissue, normal-adjacent tissue and matched blood). (**D**) UMAP of neutrophils from blood (BN) and tumor tissue (TAN) analyzed by scRNA-seq. (**E**) Volcano plot of the gene expression from BNs and TANs. (**F**) Expression of *OLR1*, *PLAU*, *CXCR2*, and *VEGFA* in BNs and TANs. (**G**) Selected multiplex immunofluorescence (mpIF) staining of LOX-1 (pink), pan-cytokeratin (turquoise), and DAPI (blue) in tumor tissue and matched normal-adjacent colorectal cancer (CRC) tissue (left panels). LOX-1 pseudo-DAB staining and DAPI as hematoxylin in tumor and matched normal-adjacent colorectal cancer (CRC) tissue (right panels). Scale bar: 100 μm. See also Figure S4A. (**H**) Top 10 markers for BN1-BN3 and TAN1-TAN4 sub-clusters. The marker gene quality is reflected by the area under the receiver operator characteristics curve (AUROC; 1 = marker gene perfectly distinguishes the respective cluster from other clusters in all patients; AUROC 0.5 = no better than random). (**I**) Partition-based graph abstraction (PAGA) based on RNA velocities, projected on the UMAP plot of neutrophils from blood and tumor tissue. Edges represent cell-type transitions called by PAGA. (**J**) Expression of *HLA-DRA* in neutrophils from blood and tumor tissue, analyzed by scRNA-seq. **K**) Quantification of HLA-DR expression on neutrophils by flow cytometry across tumor tissue, normal-adjacent tissue and blood. Each dot represents the mean value for an individual patient (n = 12). (**L**) Intratumoral HLA-DR^+^ neutrophils and survival in early stage CRC patients.

We observed distinct transcriptomic differences between blood *versus* tumor-associated neutrophils (TAN) (Figure 4D) with 2,214 differentially expressed genes (padj<0.05; Figure 4E). We evaluated previously described marker genes distinguishing TAN from normal tissue neutrophils (NAN) across cancers, including *OLR1*, *VEGFA*, *CXCR2*, and *PLAU*^30^. TAN markers (*OLR1*, *PLAU*, *VEGFA*) were highly upregulated in TAN of CRC patients, whereas *CXCR2*, a marker of mature neutrophils, was highly expressed in blood neutrophils but downregulated in TAN (Figure 4F). Spatial analysis of Lectin-type oxidized LDL receptor 1 (LOX-1; encoded by *OLR1*) expression using multiplex immunofluorescence (IF) staining demonstrated the presence of segregated LOX-1⁺ neutrophil cell clusters exclusively in CRC tumor tissue, but not in normal-adjacent colonic tissue (Figure 4G and S4A). Of note, LOX-1 has previously been identified as a marker for TAN in lung cancer^15^, indicating LOX-1 as a putative tumor-agnostic TAN marker.

Sub-clustering of neutrophils revealed three distinct blood neutrophil clusters (BN1-BN3) and four tumor-associated neutrophil clusters (TAN1-TAN4) (Figure 4H). Trajectory analysis indicated a progressive transition from BN1 to BN3 and subsequently to TAN1 (Figure 4I), demonstrating that neutrophil migration from the bloodstream into tissue is accompanied by distinct transcriptomic changes. From TAN1, neutrophils diverged into two trajectories: one leading to a putative pro-tumor TAN phenotype (TAN4) characterized by high *IL1A* expression, a key driver of emergency myelopoiesis that fuels inflammation and immunosuppression in the TME^31^, and another toward a phenotype defined by high expression of antigen-presentation-associated markers, such as *HLA-DRA* (TAN2/TAN3) (Figure 4J). We then validated HLA-DR protein expression on neutrophils using flow cytometry and confirmed that the expression was markedly higher in TAN compared to both blood neutrophils and NAN (Figure 4K). Neutrophils with antigen-presenting capabilities, referred to as hybrid neutrophils^30^, have previously been identified in early-stage lung cancer and are associated with an anti-tumor phenotype^15,32^. The analyses of the expression data showed that in these neutrophil subtype several HLA-DR genes were upregulated whereas calprotectin (S100A8/S100A9) was downregulated (Figure S4B). Analysis of the cell-cell communication using eight ligand-receptor methods (LIANA+)^33^ indicated HLA-DR^+^ neutrophil-T cell interactions (Figure S4C).

To investigate the prognostic value of HLA-DRA^+^ neutrophils we used tissue microarrays constructed from an independent patient cohort (cohort II, n=210) and multiplexed IF stainings (Figure S4D and S4E). HLA-DRA^+^ neutrophils were detectable in both tumor core and the invasive margin as clusters of cells, albeit at different fractions (8% and 7% of the total neutrophils, respectively, Figure 4SF). Patients with tumors with high densities of HLA-DRA^+^ neutrophils in the tumor core had better outcome compared to patients with low densities (Figure 4L). This effect was stronger for early stages compared to more advanced stages (Figure S4G). Additionally, the presence of HLA-DRA^+^ neutrophils in the tumor margin had an opposite effect in both, early and late stage tumors (Figure S4G), suggesting that the localization of these cell types plays a major role in the control of tumor progression.

### Spatial single-cell multi-modal profiling reveals functional neutrophil niches

Our data showed that neutrophils in the TME occupy spatially distinct areas or patches (Figure 4G and Figure S4D), suggesting the existence of specific cellular compositions and discrete histological properties: so-called cellular neighborhoods^23^ or cell niches. However, due to the limited number of features in the used multiplexed IF assay, it was not possible to deduce cell-cell interactions from these spatial profiling dataset. In order to quantify spatial niches in general, and specifically neutrophil niches we therefore applied spatial transcriptomic technique with 380 genes and cellular resolution (10x Genomics Xenium platform), and used FFPE tumor samples from CRC patients (cohort III, n=15). In total, 3.7 million cells were profiled resulting in 217 million expression values. The spatial single-cell transcriptomic profiling revealed distinct spatial architecture and cell type composition in both, center of the tumor and the invasive margin (Figure 5A). Whereas normal adjacent tissue sections recapitulated the known morphology of healthy colon epithelium, tumor cores and invasive margins were more heterogeneous in their spatial architectures and immune infiltration patterns. The cellular composition of the samples was heterogeneous and neutrophils were detectable in all samples (Figure 5A). Neutrophils were spatially localized as aggregates or clusters in both the tumor core and the invasive margin (Figure S5A). This spatial organization of neutrophils was confirmed using H&E stainings from the same slide as well as immunohistochemistry (IHC) stainings of consecutive FFPE slices confirmed by two board certified pathologists (S.O. and M.G) (Figure S5A).

**Figure 5.**
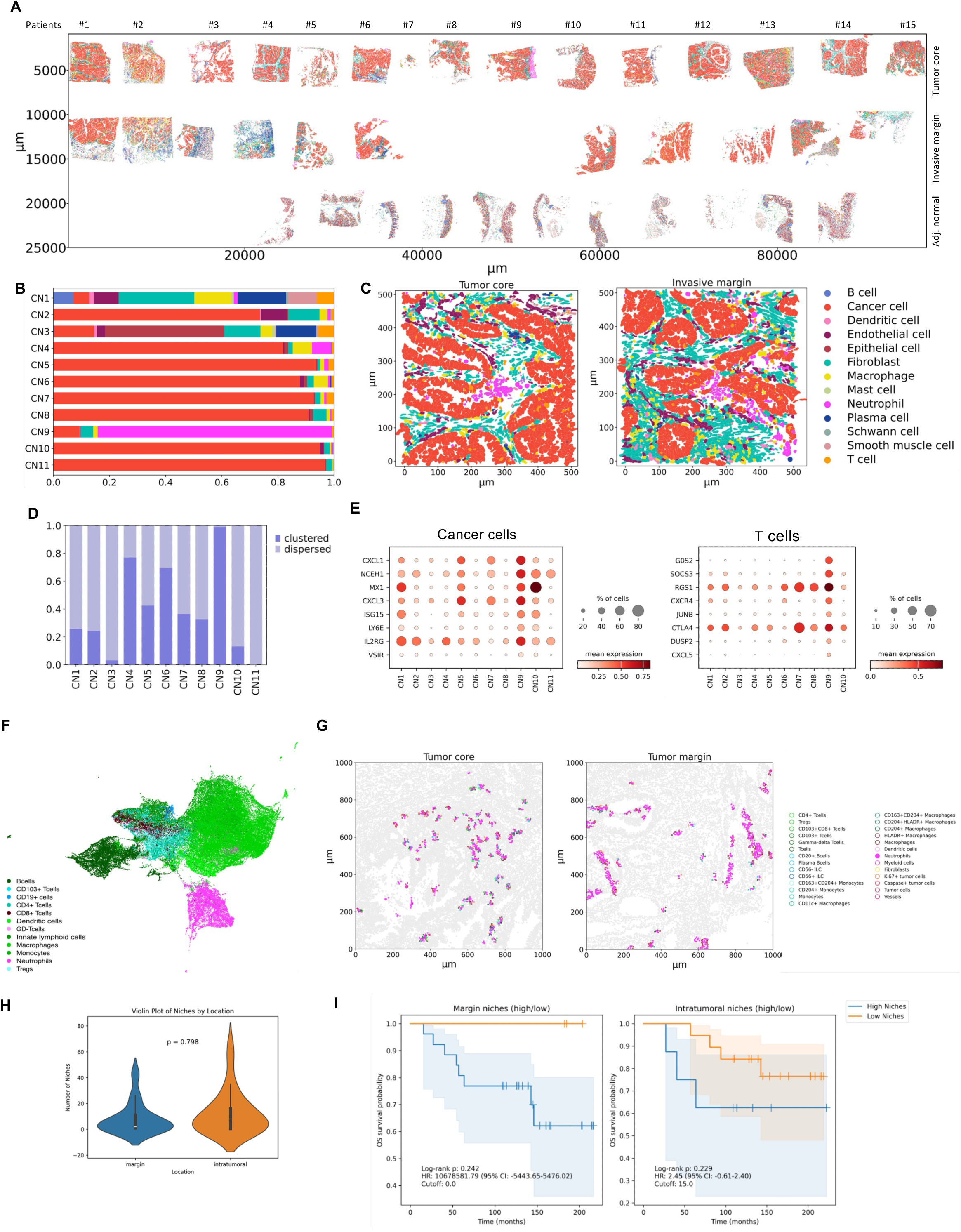
Spatial single-cell multi-modal profiling. (**A**) Cell types identified from imaging-based spatial transcriptomics profiling (10X Xenium platform) of 37 CRC tumor sections (tumor cores, invasive margins and adjacent normal tissue from 15 patients). (**B**) Cell niches quantified using the tool NicheCompass. (**C**) Representative image of neutrophil clusters located in the tumor core and invasive margin. See also Figure S5A. (**D**) Neutrophil clusters in each niche. (**E**) Top 8 genes upregulated in cancer cells and T cells within CN9. (**F**) Uniform manifold approximation and projection (UMAP) of imaging mass-cytometry profiling data from tumors from CRC patients (n = 29). (**G**) Neutrophil clusters identified using the imaging mass-cytometry data at the tumor core and invasive margin. (**H**) Violin plot of the number of neutrophil niches (= clusters) in each patient stratified by location (margin, intratumoral). (**I**) Kaplan-Meier curves for the neutrophil niches (= clusters) identified at the tumor core and invasive margin. Optimal cutoff values were determined to define low and high neutrophil niches, while minimizing the p-value.

We then aimed to quantitatively define cellular niches from their biological functions and used a graph deep-learning method that utilizes existing knowledge of inter- and intracellular interaction pathways to identify such functional cellular niches (Niche Compass)^34^. At the resolution level of two niches, we identified cancer and non-cancer niches, whereas at the resolution of three niches we identified cancer, boundary, and stroma niches (data not shown). At the next resolution level we quantified eight cancer cell-enriched cell niches (CN2, CN4-CN8, and CN10-11), one fibroblast-enriched niche (CN1), one normal epithelial cell-enriched niche (CN3) and one neutrophil-enriched niche (Figure 5A and B and Figure S5B). The cancer cell-enriched cell niches had different fractions of infiltrating immune cells and in nine niches there were neutrophils with varying cellular fractions (Figure 5B). One cellular niche (CN9) was dominated by neutrophils which were clustered in both the tumor core and the invasive margin (Figure 5C). Neutrophils were clustered also in other cellular niches, albeit at different fractions of the total neutrophil count in the specific niche (Figure 5D). The cellular niche with the largest fraction of neutrophils (CN9) included cancer cells, fibroblasts, macrophages, and T cells. The analysis of the gene expression of the cancer cells within the neutrophil niche CN9 revealed elevated expression of the chemokines *CXCL1*, and *CXCL3* (Figure 5E). Analysis of the gene expression of the T cells in the neutrophil niche CN9 revealed elevated expression of chemokine receptor *CXCR4* (Figure 5E). The identified chemokines likely play a role in driving cancer-cell mediated localized neutrophil aggregation and cluster formation. Hence, our cell-cell interaction analyses of the CRC TME (Figure S5C) provided evidence for communities of cells coordinating specific function within a tissue.

To validate the findings based on mRNA expression in single cells we then used an orthogonal protein-based method (imaging mass-cytometry (IMC)) and profiled FFPE samples from patients with stage III colon cancer treated with adjuvant chemotherapy (CAPOX) and for whom relapse and overall survival data were available (cohort IV, n=29). We previously developed and evaluated a panel of 41 antibodies for IMC on FFPE samples for a comprehensive overview of the TME and cancer-immune cell interactions^35^, including lineage and functional immune cell markers, surrogates of cancer cell states (proliferation, apoptosis) and structural markers (epithelium, stroma, vessels). Following data-driven identification of single-cell phenotypes segmentation, and image analysis, we quantified the densities of major cell types (Figure 5F). The spatial protein profiling revealed similar cellular composition of the TME as identified using single-cell mRNA profiling (Figure 5F). Given the limited number of features of the IMC assay, quantification of functional cell niches based on protein expression data was not possible. We therefore used a cell neighborhood analysis to define niches by defining cell nuclei and associating polygons (Voronoi diagram) to each nucleus^35^ (see Methods), thereby allowing cells of different sizes and distances to be assessed as neighbors. We used a permutation approach^35^ to identify pairwise interactions between cell phenotypes that occurred more or less frequently than expected by chance (see Suppl. Table S4). The analysis revealed neutrophil clusters similar to the clusters quantified using transcriptomic data (Figure 5G). It appears that the neutrophil clusters are more frequent in the tumor center compared to the invasive margin, albeit the differences in this cohort were not significant (Figure 5H). Analyses of the survival data, employing an optimized cutoff that minimizes the p-value, showed that the neutrophil niches in both regions are associated with worse prognosis (Figure 5I).

Overall, the spatial single-cell transcriptomic profiling of the TME revealed cellular niches composed of different fractions of cancer cells, immune cells, endothelial cells, and fibroblasts, with distinct cell-cell communication patterns. Spatial single-cell proteomics profiling in a larger independent patient cohort confirmed the spatial organization of neutrophils in distinct niches in both the tumor core and the invasive margin.

### Mouse neutrophils recapitulate human neutrophil phenotypes in various compartments

Our single-cell gene expression data indicated changes of the neutrophil phenotype following migration from the bloodstream into tumors. In order to investigate the neutrophil phenotype changes along the bone marrow-blood-tumor axis we used an orthotopic mouse model based on tumor organoid transplantation (Figure S6A). AKPS organoids – harboring the four most common CRC-driving mutations (*Apc*-LOF (loss of function), *p53*-LOF, *Kras*-GOF (gain of function), *Smad4*-LOF) – were successfully engrafted following colonoscopy-guided injection into the colonic mucosa of C57BL/6JRj mice^36^. Tumors developed within six to eight weeks and progressively expanded through the colonic layers toward the peritoneal cavity (Figure S6A), mirroring some of the invasive characteristics of late-stage human CRC. Notably, neutrophil infiltration was significantly higher in the tumor compared to the healthy colon (Figure 6A-C). Interestingly, there was already an increase of neutrophils within the blood of AKPS-tumor bearing compared to naive mice (Figure 6C) suggesting a systemic effect during tumor development. Analysis of neutrophil activation markers like Siglec-F, CD11b, CD80 and PD-L1 further showed a progressive activation of neutrophils from the blood to the tumor (Figure 6D-G) while MHC-II proteins were upregulated in tumor and colon tissue compared to blood neutrophils (Figure 6H). In contrast, only little phenotypic changes were observed in the bone marrow (Figure S6D). Overall, the results confirm the validity of the orthotopic mouse model for human CRC.

**Figure 6.**
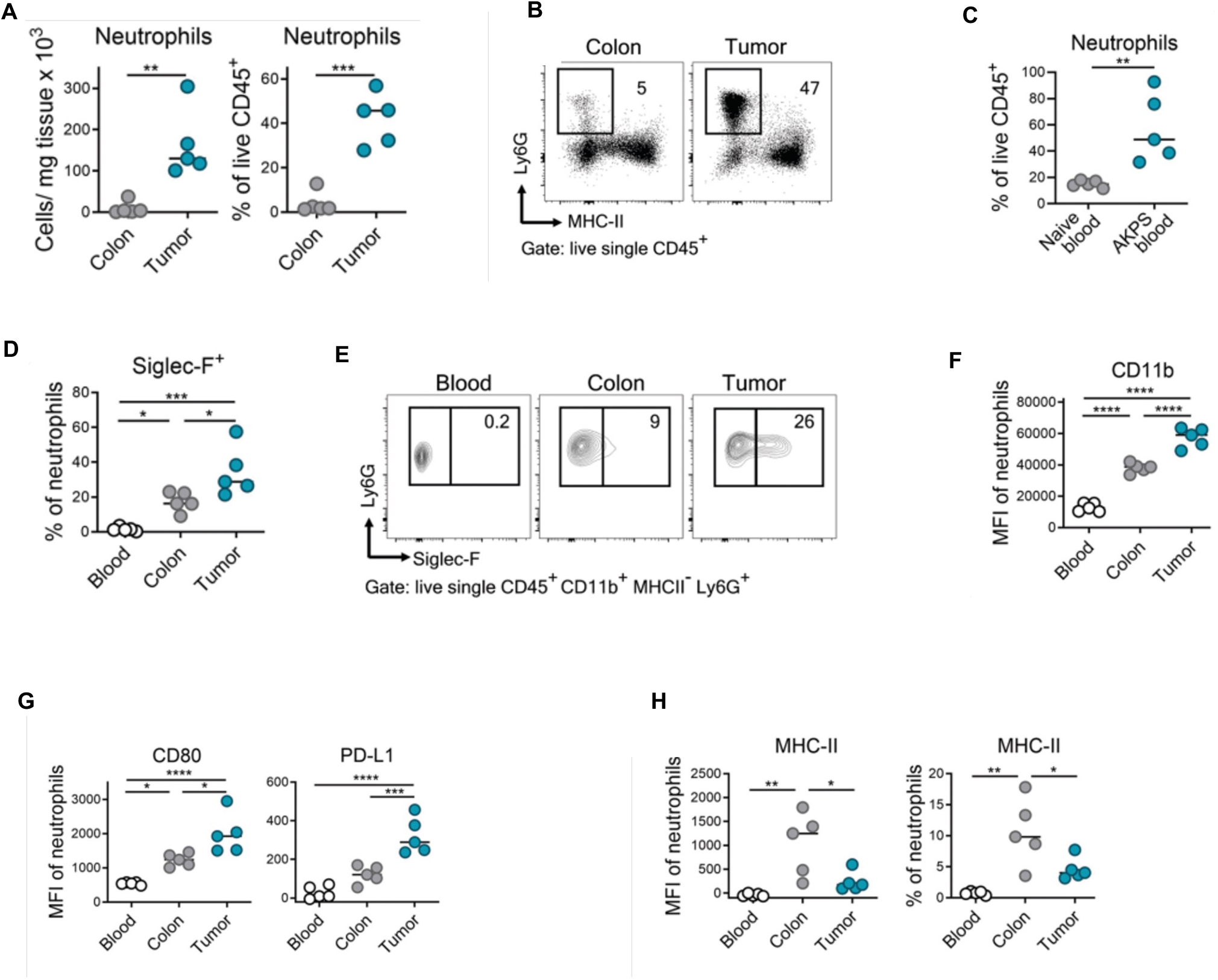
Neutrophils in a murine orthotopic organoid transplant model for CRC. Mouse AKPS tumor organoid cells harbouring the four most common CRC driver mutations (*APC*^KO^, *KRAS*^G12D^, *P*53^KO^, *SMAD*4^KO^) were injected intra-rectally into 8- to 10-week-old C57BL/6J males. At the study endpoint (4-6-weeks post-injection), blood, tumor and adjacent colonic tissue were harvested for flow cytometric analysis. (**A**) Neutrophil numbers per milligram tissue (left) and frequency among live CD45^+^ leukocytes (right). (**B**) Representative plots of neutrophil frequencies among CD45^+^ cells, as determined by flow cytometry. (**C**) Neutrophil frequencies among CD45^+^ cells in the blood of naïve versus AKPS-injected mice. (**D**) Frequencies and (**E**) representative flow-cytometry plots of Siglec-F-expressing neutrophils among total neutrophils in blood, colon and tumour of AKPS-bearing mice. Mean Fluorescence Intensity (MFI) of cell-surface CD11b integrin (**F**) or CD80 and PDL1 (**G)** expression in neutrophils. (**H**) MFI (left) and frequencies of MHC II expression among neutrophils. *, P < 0.05; **, P < 0.01; ***, P < 0.001; ****, P < 0.0001; as calculated by two-tailed unpaired Student’s t-test for comparison of two groups (**D, F**) or by one-way ANOVA for statistical comparison of more than two groups (**A-H**). n = 5, medians are displayed.

### Cross-species analyses reveal tumor-induced granulopoiesis in CRC

We then performed scRNA-seq using CD45-enriched leukocytes isolated from mice six weeks post-AKPS-injection. Colons and blood of naïve (control) mice were further included as controls. Additionally, CD45-enriched leukocytes were isolated from the bone-marrow of both tumor and non-tumor bearing mice. Neutrophils from all tissues and phenotypes were extracted and integrated (Figures 7A and B). Clustering analysis identified 10 clusters with distinct transcriptional patterns (Figure 7C) and a high concordance of BN and TAN marker genes derived from our human neutrophil dataset (Figure S7A), especially within the colon and tumor specific clusters. We then performed pseudotime trajectory analysis, which showed a progressive transition of neutrophils from the bone-marrow to the blood and colonic tissues (Figures 7D-F). Differential gene expression (DEG) analysis between tumor and healthy bone-marrow and blood revealed that neutrophils already changed their transcriptional phenotype prior to their recruitment into the tumor tissue (Figures 7G and H), indicating tumor-induced granulopoiesis. Gene set enrichment analysis (GSEA) of KEGG pathways showed changes in the metabolic phenotypes (Figure S7B). Overall, the results from the single-cell profiling suggest tumor-induced granulopoiesis which is likely driven by secreted factors^37–40^.

**Figure 7.**
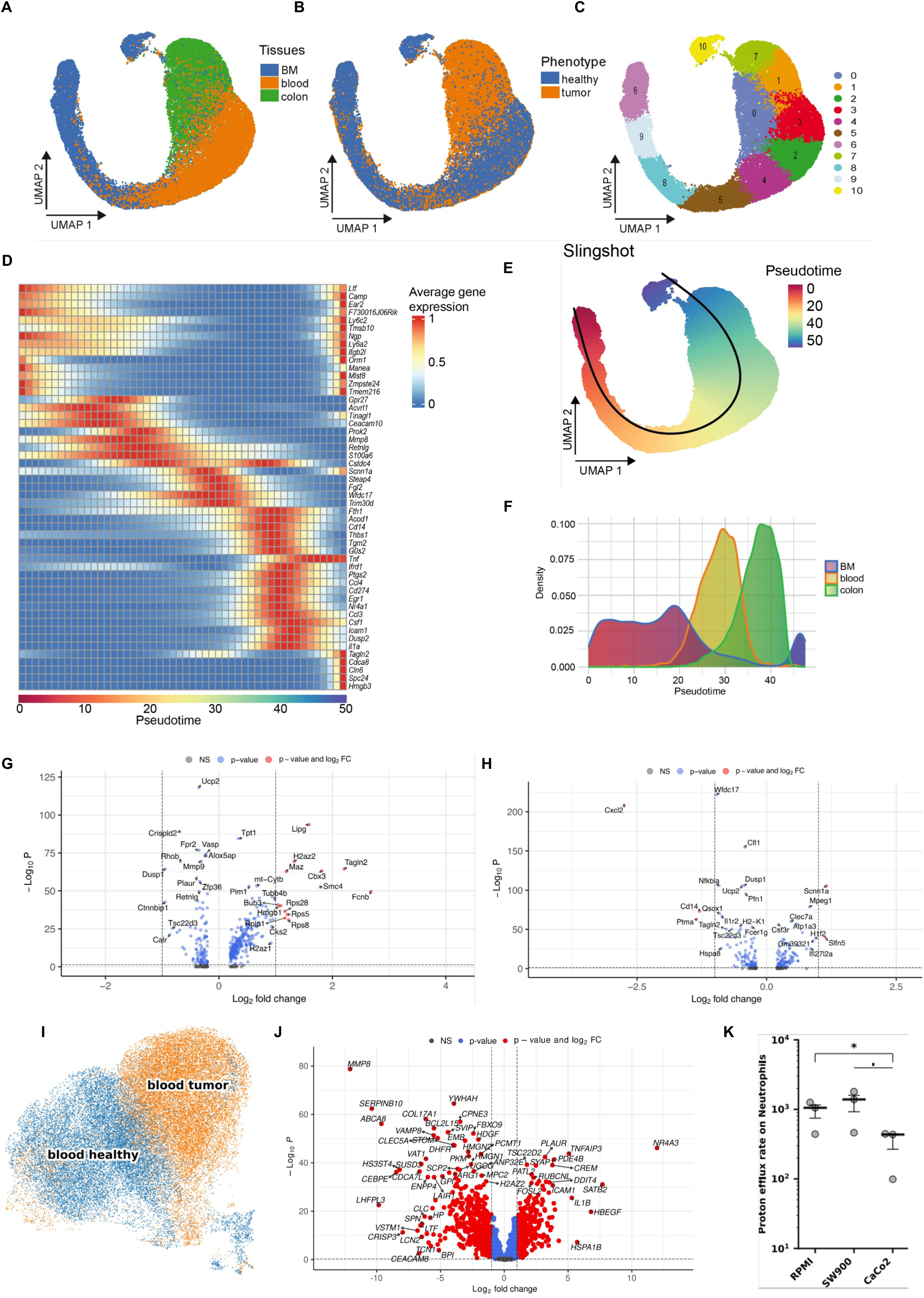
Neutrophil phenotypes along the bone marrow-tumor axis. Uniform manifold approximation and projection (UMAP) of integrated neutrophils from tumor, colon, blood and bone marrow of AKPS tumor bearing and non-tumor bearing mice. Cells are colored by (**A**) tissue type (colon is a combination of colon, adjacent colon, tumor, and disseminated tumor), (**B**) phenotype and (**C**) clusters generated via Louvain algorithm with multilevel refinement. (**D**) Heatmap depicting the average gene expression of selected genes along the pseudotemporal trajectory (Adjusted p-values ≤ 0.05, Wald-text, FDR correction). (**E**) UMAP representing pseudotime from inferred trajectory using Slingshot. (**F**) Density plot representing neutrophils from different tissues ordered by pseudotime. Volcano plots of differentially expressed genes from **(G)** bone-marrow and (**H**) blood neutrophils between healthy and tumor-bearing mice (Wilcoxon Rank-Sum test, Bonferroni corrected p-values). (**I**) UMAP of blood neutrophils from CRC patients and controls. (**J**) Volcano plot of gene expression data from blood neutrophils from CRC patients and healthy individuals. (**K**) Proton efflux rate (PER), as an indicator of glycolysis, of human neutrophils activated with PMA, after 24 hours culture with SW900 supernatant, CaCo2 supernatant or RPMI control medium. The data shown are the mean and SD, tested for differences using paired Student’s t test, n = 3 biological replicates. **(A-H)** n = pool of 3 (non-tumor blood and bone marrow) or 5 (non-tumor colon and tissues from AKPS tumor-bearing mice) mice.

To provide further evidence for tumor-induced granulopoiesis in patients we carried out scRNA-seq profiling using blood samples from healthy controls and compared the expression profiles of neutrophils with the expression profiles of blood neutrophils from the CRC patients (Figure 7I). The results showed profound changes in the expression with 2287 genes differentially expressed (padj<0.05; Figure 7J). GSEA on KEGG pathways indicated changes in the metabolic phenotypes of the blood neutrophils from CRC patients (Figure S7C). We then carried out functional experiments by quantification of neutrophil activation using Seahorse technology. Neutrophils isolated from blood of healthy donors were cultured with supernatants from a colorectal adenocarcinoma cell line (CaCo2) and a non-small cell lung cancer cell line (SW900) for 24 hours. Upon stimulation with Phorbol 12-myristate 13-acetate (PMA) the NOX2 enzyme is activated which results in an oxidative burst, consuming a large amount of oxygen and leading to a rapid cellular metabolic shift towards glycolysis. The proton efflux rate (PER) and the oxygen consumption rate (OCR) were measured after 24 hours treatment with the supernatants. Interestingly Seahorse analysis showed a significant decrease upon stimulation in PER and OCR when neutrophils were treated with supernatant from the CaCo2 cell line compared to the control (Figure 7K and Figure S7D). This observation of changes of non-mitochondrial PER and OCR after neutrophil activation indicates context-dependent metabolic rewiring of the neutrophils.

Together, we charted for the first time neutrophil phenotypes at the single-cell level along the bone marrow-tumor axis in a mouse model of CRC. Importantly, the results provide evidence for tumor-induced granulopoiesis and metabolic rewiring of neutrophils, suggesting therapeutic strategies targeting the bone marrow.

## DISCUSSION

We built a large-scale atlas of single-cell transcriptomes of CRC through integration of 4.27 million cells from 650 patients and 1,670 samples representing 7 billion expression values. We complemented the atlas with single-cell data from four CRC cohorts with 266 patients, and enriched the atlas with immune cells with low-mRNA content including neutrophils, as well as with multi-modal spatial single-cell profiles of the TME comprising transcriptional profiles from 3.7 million cells, and protein profiles from 0.7 million cells. The exploitation of our unique big data resource led to several biological discoveries and has important translational relevance.

First, we provide a high-resolution view of the TME in CRC with 61 major cell types/states and show different cell type composition patterns. The single-cell composition of the TME in CRC enabled refined tumor classification and patient stratification into four tumor immune phenotypes: immune deserted, myeloid, B cell, and T cell subtypes. These findings may have important implications for the development of new cancer immunotherapy strategies in CRC. For example, combination therapies that target both, myeloid cells and lymphoid cells could represent an immunotherapeutic approach to treat M subtypes as shown recently in a melanoma mouse model^41^. Similarly, given the heterogeneity of the intratumoral B cells and the importance of tertiary lymphoid structures^42^, further analysis of the B cell subtypes might open promising therapeutic avenues by targeting B cells.

Second, we characterized for the first time neutrophil subsets in various compartments in CRC patients and show remarkable diversity and plasticity. Of particular interest for cancer immunotherapy is the HLA-DRA^+^ TAN phenotype with antigen-presenting feature that is associated with better survival outcome. This observation might explain the conflicting reports on neutrophils suggesting their dual function as pro-tumorigenic and anti-tumorigenic cells. Such conversion of neutrophils to antigen presenting cells may elicit anti-tumor immunity as recently shown in a murine model^43^. We show the heterogeneity of short-lived neutrophil populations in blood and long-lasting neutrophil populations in colorectal cancer tissue on transcriptional level (Figure 4H), which is a characteristic for a chronic inflamed microenvironment^22^. Noteworthy, we also provide evidence for genotype-immune phenotype relationships and show that the enrichment or depletion of neutrophils in the TME is associated with specific tumor genotypes.

Third, single-cell spatial transcriptional profiling revealed that cells in the tumor core and the invasive margin build functional cellular niches with different cellular compositions including several cancer cell-enriched niches, fibroblast-enriched niche, normal epithelial cell-enriched niche, and one neutrophil-enriched niche. The functional consequence of these spatial architecture could not be elucidated due to the limited coverage of intra- and extra-cellular pathways of the used probes. Additional studies using larger number of probes will be necessary to provide insights into this cellular architecture. Of particular interest is a neutrophil-enriched niche in which neutrophils build clusters of cells. Within this niche we observe intracellular cell-cell interactions of neutrophils with cancer cells, fibroblasts, macrophages, and T cells. The driving force for this type of myeloid structures remains unknown and encourages further studies in the context of CRC and possibly other solid cancers.

Finally, using an orthotopic mouse model and cross-species analyses, we show that tumor-derived signals modulate the neutrophils in the bone-marrow, implicating tumor-induced granulopoiesis. Expression and functional data indicated metabolic rewiring in the bone marrow and suggest imprinting the neutrophil cell fate via progenitors. Thus, reprogramming neutrophils into anti-tumorigenic states for more effective therapies of cancer^43^ requires targeting of the bone marrow. Such an approach to specifically modulate the innate immune system by inducing trained immunity using bone marrow-avid nanobiologics and elicit anti-tumor response was recently developed^44^.

Beyond these biological insights, the results from this study have also important translational implications. As shown here, TANs can acquire antigen presenting properties and such conversion of abundant neutrophils to antigen-presenting cells could overcome the limitations of the low abundance of cross-presenting DCs. Additionally, our study highlights CRC as a systemic disease that requires therapeutic strategies beyond the primary disease site. Such a strategy was recently shown for lung cancer^45^. However, as we demonstrate here, the disease context has profound effects on the neutrophil phenotypes and hence, requires further disease-specific investigations.

In conclusion, we provide a CRC atlas with single-cell resolution which is publicly accessible and interactively explorable through cell-x-gene (https://doi.org/10.1101/2021.04.05.438318) (https://crc.icbi.at) that allows visualization of metadata and gene expression. The biological insights we present here and future discoveries arising from the exploitation of the high-resolution CRC atlas have the potential to provide the basis for developing combination therapies for CRC patients who are not sufficiently responding to immune checkpoint blockers.

## ACKNOWLEDGMENTS

The authors would like to thank Martina Reschreiter, Anna Mair, Fabienne Isabelle Nocera, Vera Danklmaier, Annabella Pittl, Corinna Antonia Griesbaum, Petra Schumacher, Elisabeth Hoflehner and Manuel Trebo for their excellent technical support. This work was supported by the European Research Council (grant agreement No 786295 to ZT), by the Austrian Science Fund (FWF) (project I3978 to ZT). ZT is a member of the German Research Foundation (DFG) project TRR 241(INF). VM was funded by the Austrian Science Fund (FWF) 10.55776/DOC82. ICA was funded through an Eccellenza Professorial Fellowship from the Swiss National Science Foundation PCEFP3_187021 and through a Tandem Grant (2022 call) of the ISREC Foundation. NB was funded by resources of the State of Tyrol F.47895/6-2023. The results published here are in part based upon data generated by the TCGA Research Network: https://www.cancer.gov/tcga.

## METHODS

### Patient cohorts and recruitment

Experimental data was generated using samples from four CRC patient cohorts. Patient samples from cohort I (n = 12) were used for single-cell RNA-seq. Tumor and matched adjacent normal tissues from CRC patients who underwent surgical resections at the Department of Surgery, Medical University of Innsbruck, Austria were obtained. Additionally, autologous whole blood samples were collected during surgery. Prior to study enrollment, all patients signed the informed consent for participation in the study that was approved by the ethical committee of the Medical University of Innsbruck (AN2014-0282 342/4.3 425/AM3 (4516a)). Patient samples from cohort II (n = 210) were used for mIF analysis. Samples and patient data were obtained from the University Cancer Center Frankfurt (UCT) in collaboration with the Senckenberg BioBank at the University Hospital Frankfurt. Written informed consent was obtained from all patients and the study was approved by the institutional Review Boards of the UCT and the Ethical Committee at the University Hospital Frankfurt (project-number: SGI-11-2018). FFPE samples from cohort III (n = 15) were used for profiling using spatial transcriptomics. The study was approved by the ethical committee of the Medical University of Innsbruck (AN2014-0282 342/4.3 425/AM3 (4516a)). Patient samples from cohort IV (n = 29) were analyzed using IMC. These patients uniformly had R0-resected stage III colon cancer and received adjuvant CAPOX treatment. This analysis was approved by the Ethics Committee of the Provincial Government of Salzburg, Austria (415-E/2343/5-2018). Patients were otherwise unselected and treated at the Department of Internal Medicine III, Paracelsus Medical University, Salzburg, Austria.

### Dissociation of normal and tumor colorectal tissue

Surgically resected human colorectal tumor and adjacent normal tissues were enzymatically and mechanically dissociated into single cells as described in detail in the following publication^46^. Briefly, after supervised sampling of tumor and adjacent normal CRC tissues from a pathologist, the specimens were transported to the lab on ice, washed, weighed and minced into small pieces (<1 mm). The tissue pieces were further enzymatically and mechanically digested by using DMEM medium (Gibco) with Liberase DH (Roche) and DNAse I (Roche/Sigma-Aldrich) on a gentleMACS dissociator with heaters (Miltenyi Biotec) at 37°C for ∼45 min (selected program: 37C_h_TDK_1). The obtained single cell solutions were sieved through a 100 µm cell strainer (pluriSelect) and kept on ice.

### BD Rhapsody library preparation and sequencing

Freshly isolated single-cells were immediately processed with the BD Rhapsody scRNA-seq platform (BD Biosciences). Red blood cells were removed from dissociated tissues and from whole blood samples using the BD Pharm Lyse lysing solution (BD Biosciences). Cell number and viability were assessed with the BD Rhapsody scRNA-seq platform using Calcein-AM (Invitrogen) and Draq7 (BD Biosciences). The BD Single-Cell Multiplexing Kit (BD Biosciences) was used to combine and load multiple samples (human colorectal tumor tissue, adjacent normal tissue, blood) onto a single BD Rhapsody™ cartridge (BD Biosciences). Sample-tag staining was performed according to the manufacturer’s protocol (sample-tag staining at room temperature for 20 min and washing by centrifugation at 400 g for 5 min). Single-cell isolation in microwells (cell load: 20 min incubation at room temperature, 20°C ± 2°C) with subsequent cell-lysis and capturing of poly-adenylated mRNA molecules with barcoded, magnetic capture-beads was performed according to the manufacturer’s instructions. Beads were magnetically retrieved from the microwells, pooled into a single tube before reverse transcription. Unique molecular identifiers (UMIs) were added to the cDNA molecules during cDNA synthesis. Whole transcriptome amplification (WTA) and sample-tag sequencing libraries were generated according to the BD Rhapsody single-cell whole-transcriptome amplification workflow. The quantity and quality of the sequencing libraries was analyzed with the Qubit dsDNA HS (High Sensitivity) assay kit (Invitrogen) and the 4200 TapeStation (Agilent) system. Libraries were sequenced on the Novaseq 6000 system (Illumina) targeting a sequencing depth of 50.000 reads/cell.

### Generation of the atlas

The single-cell atlas was compiled from publicly available datasets^47–93^, including data from previously published colorectal cancer (CRC) studies, our own data (cohort I), and data from 12 additional non-CRC studies for control purposes, including only healthy, non inflamed control colon samples from adult individuals (see Suppl. Table S1). These studies, published between May 2017 and July 2024, utilized six different sequencing platforms, including the widely used 10x Chromium (10x Genomics) as well as Smart-seq2^94^, SMARTer (C1), scTrio-seq2, GEXSCOPE (Singleron), TruDrop^95^ and DNBelab C4^96^. Our own data was generated using the microwell-based BD Rhapsody scRNA-seq platform (BD Biosciences). We carefully chose studies that followed comparable protocols for sample processing and data generation, such as sequencing of whole cells, and had raw gene expression data as well as cell-level metadata available to map cells to sample information. Studies using flow cytometry-based cell sorting prior to sequencing were not excluded, as they offer important information about rare cell types^97,98,99^. To distinguish between normal colon samples from CRC patients and healthy individuals, we use the terms “healthy normal” for samples from healthy individuals and “adjacent normal” for colon samples near tumor tissue.

### Preprocessing and quality control of scRNA-seq data

We distinguish between studies, which are scientific publications, and datasets, which consist of scRNA-seq samples processed with the same sample preparation and experimental platform. A study can include one or more datasets. Fastq files of our cohort I dataset were processed using the Seven Bridges -BD Rhapsody™ WTA Analysis Pipeline version 2.0. In addition, fastq files were available for samples from the studies Bian_2018_Science, Che_2021_Cell_Discov, Conde_2022_Science, Elmentaite_2021_Nature, GarridoTrigo_2023_Nat_Commun, Guo_2022_JCI_Insight, He_2020_Ge nome_Biol, James_2020_Nat_Immunol, Ji_2024_Cancer_Lett, Joanito_2022_Nat_Genet, Lee_2020_ Nat_Genet, Wang_2020_J_Exp_Med, Wang_2021_Adv_Sci, Yang_2023_Front_Oncol and were obtained from the identifiers specified in the key resource table. Smart-seq2 data was processed using the nf-core RNA-seq pipeline v3.12.0 (https://doi.org/10.5281/zenodo.7998767), 10x datasets were processed with cellranger v7.1.0 (10x Genomics) using the nf-core scRNA-seq pipeline v2.4.1^100^ (https://doi.org/10.5281/zenodo.8399014) with the GRCh38 reference genome and GENCODE v44 annotations. All other datasets were obtained as count tables from their respective identifiers. All raw count matrices were loaded into AnnData^101^ and further processed with scverse tools^102^. For samples with available fastq files, ambient RNA was removed using scAR^103^ as implemented in scvi-tools^104^. Quality control was performed using scanpy^105^ by retaining cells with (1) >400 transcripts, (2) >100 genes, and (3) <50 % mitochondrial transcripts, followed by 3 MADs (median absolute deviations) filtering on individual samples^106^.

### Integration of scRNA-seq datasets

Individual datasets were merged into a single AnnData object. Since genome annotations partly differed between the datasets, we mapped gene symbols to the ensembl ids of the used annotation version and merged on stable ensembl ids. If gene ids were missing from a dataset, the values were filled with zeros. Genes that were missing in more than 19 datasets (25%) were excluded altogether. We integrated the datasets using the scANVI algorithm^107,104^, as it has been demonstrated to be one of the top-performing methods for atlas-level integration and to scale to >1M cells^108^. Given that scANVI requires cell-type annotations for one input dataset, we manually annotated the major sample types, including tumor and polyp, normal, metastasis, blood, and lymph nodes, as these were expected to represent the main sources of biological variation. These five “seed” datasets were annotated based on unsupervised clustering as described below. Raw counts were used as input for scANVI. Counts from plate-based assays such as Smart-seq2 were scaled by the gene length as recommended on the scvi-tools website. The scANVI model was initialized with a pre-trained scVI model^109^, as recommended in the scvi-tools tutorial. The scVI model was trained on the 6000 most highly variable genes as determined with scanpy’s^105^ *pp.highly_variable_genes* with parameters *flavor=”seurat_v3”* and *batch_key=”dataset”*. Each sample was considered as an individual batch for both scVI and scANVI. Other than that the algorithms were run with default parameters.

### Unsupervised clustering and cell-type annotation

We computed UMAP embeddings^110^ and unsupervised Leiden-clustering^111^ with scanpy^105^, based on a cell-cell neighborhood graph derived from scANVI latent space. Coarse, lineage-specific clusters were iteratively sub-clustered to identify cell-types at a more fine-grained resolution. Cell type clusters were annotated based on previously reported marker genes^54^.

### Patient stratification

We stratified patients into immune phenotypes based on immune cell-type fractions. We selected all patients with primary tumor samples and excluded enriched datasets. Neutrophil fractions were excluded, since they are not appropriately captured in the majority of datasets. Cell-type fractions of primary tumor samples were loaded into a patient × cell-type AnnData container. Dataset-specific batch-effects were removed using a linear model as implemented in *scanpy.pp.regress_out*. Patients were clustered using graph-based Leiden clustering with the “correlation” distance metric for computing the neighborhood graph. Patient clusters were labeled according to their predominant cell-types.

### Differential gene expression testing

We used DESeq2^112^ on pseudo-bulk samples for differential expression testing which has been demonstrated to perform well and properly correct for false discoveries^113^. For each cell type and sample, we summed up transcript counts for each gene that is expressed in at least 5% of cells using decoupler-py^114^. Pseudo-bulk samples consisting of fewer than 10 cells were discarded. We compared primary tumor samples from tumor *vs.* normal, tumor *vs.* blood (*sample type*), primary tumor samples from the patient subtypes M/B/T/desert (*group*), BN *vs.* TANs (*cell_type_fine*), and Neutrophil clusters (BN1-3, TAN1-4), including the patient as a covariate for paired comparisons, else we included the dataset. For comparisons between multiple groups, we used contrasts with sum-to-zero coding. P-values were adjusted for multiple hypothesis testing with independent hypothesis weighting (IHW)^115^.

### Pathway, TF and cytokine signaling signatures

We performed pathway, transcription factor (TF), and cytokine signaling analysis on primary tumor samples with PROGENy^26,116^, and CytoSig^27^, respectively. Scores were computed using the *decoupler-py*^114^ package. Transcription factor activity was inferred from the Wald statistics of the differential expression analysis output, using a multivariate linear model and the CollecTRI^117^ database. Only regulons with the highest confidence levels “A” and “B” were used. Activities were defined as significantly activated or suppressed based on an FDR threshold of less than 0.1. The top 1,000 target genes of the progeny model were used, as recommended for single-cell data. The cytosig signature matrix was obtained from the *data2intelligence/CytoSig* GitHub repository and used with the scoring function implemented in the *progeny-py* package. The methods were run with the options *num_perm=0, center=True, norm=True scale=True,* and *min_size=5.* No permutations were used, as we perform statistics in a separate step at the level of biological replicates. Pathway-, transcription factor-, and cytosig scores were then compared between *condition* (tumor *vs*. normal) and patient *group* (T *vs*. B *vs.* M *vs.* deserted) using an ordinary least-squares (OLS) linear model, as implemented in the statsmodels package^115^.

### Liana+ of scRNA-seq data

We performed liana analysis on the atlas using LIANA+^33^ combining DEA results from HLA-DR+ neutrophils. Ligand–receptor interactions were inferred using likelihood ratios (consensus, threshold 0.1) For visualization, we generated a dot plot summarizing the ligand–receptor interactions. *ligand_pvalue* reflects ligand significance in DEA, while *Interaction_stat* integrates expression levels and statistical significance.

### RNA velocity analysis

We performed RNA-velocity analysis on our own dataset (cohort I) using velocyto and scVelo. BAM files generated by the BD Rhapsody WTA analysis pipeline were preprocessed with samtools to ensure compatibility with velocyto (see preprocessing/bd_rhapsody/velocyto.nf in our git repository for more details).. The resulting loom files from velocyto were then processed in scVelo to estimate and visualize RNA velocities, following the scVelo tutorial. To analyze cellular transitions, we computed Partition-based Graph Abstraction (PAGA) using the RNA velocity graph, with neutrophil subclusters as grouping variables and the parameters minimum_spanning_tree=False and use_time_prior=True. The final output was a directed graph, where transition confidences were represented as weighted edges.

### Mapping bulk RNA-seq data in the atlas using Shears

We developed Shears, a Python-based method designed to address critical limitations of existing algorithms^119,120^ to associate single cells from scRNA-seq datasets with phenotypic or clinical information of bulk RNA-seq samples (source code and documentation are available at https://github.com/icbi-lab/shears). For example the widely used developed method Scissor^132^ has several limitations: 1) The method is computationally expensive, as demonstrated by the several thousand CPU hours required to apply it to large single-cell datasets like the CRC atlas; 2) It does not adequately account for biological replicates within the single-cell data; and 3) Despite being based on a generalized linear model it is not possible to include covariates into the regression analysis. Hence, it is not possible to determine whether the observed effects are due to confounding factors. For example, when analyzing the entire dataset for associations with a mutation that occurs preferentially in COAD over READ, it is unclear whether the ’scissor cells’ are linked to the mutation itself or the tumor type. This uncertainty extends to other variables as well, such as gender, age, microsatellite instability, or co-mutations. Additionally, Scissor results are determined relative to other cells in the single-cell RNA-seq dataset. The cells that show the strongest association with the phenotype of interest are classified as Scissor-positive or Scissor-negative while L1-regularization forces other coefficients of less important cells to be zero. As a consequence, in large datasets, Scissor results show the average of which cells were chosen as “most important” over many patients. To overcome these limitations, we developed Shears, an alternative method that enhances computational efficiency for large datasets (>1M cells) while addressing the handling of covariates and biological replicates, enabling more accurate identification of cell subpopulations from single-cell data associated with a given phenotype.

Shears utilizes a two-step approach: 1) generate a bulk x single cell matrix with cell weights assuming that bulk=weights∗single cells. This is similar to deconvolution, except we have a single-cell matrix instead of a signature matrix aggregated by cell types and can be solved with e.g. linear regression. To ensure comparability between bulk and single-cell data, quantile normalization is applied, as done by Scissor. Given the large number of features, we apply a Ridge constraint to the regression. This Ridge regression estimates weights for the following equation: B=w1*S1 + w2*S2 + … + wn*Sn. In contrast to the correlation matrix used by Scissor, by utilizing a weight matrix, we can account for the relative importance of each cell, thereby reducing the risk of falsely attributing an association between a cell and the phenotype when its effect could actually be explained by other cells. 2) Compute the importance of each cell for a given phenotype using an individual linear model per cell and fitting a model with “Phenotype ∼ cell weight + covariates” for each cell. Shears uses logistic regression for mutation data and Cox regression for overall survival. The coefficient of the model can be considered as the effect size measured for each cell, while the p value provides evidence on whether the effect size for each cell is statistically significant in explaining the phenotype. Shears accounts for biological replicates by grouping data by biological replicate (e.g., patient or sample) and cell type, ensuring at least 10 cells per group and cell type, then calculating the mean coefficient weighted by -log10(p-value) to reflect both effect size and statistical significance of the phenotype on a given cell. Significant differences are determined by comparing the fractions of shears+ and shears-cells with a paired wilcoxon test with zero_method =’’zsplit’’ as implemented in the scipy package. p-values were Benjamini-Hochberg-adjusted and considered significant at an FDR <0.01.

Shears was used to associate phenotypic data from the combined TCGA and AC-ICAM bulk RNA-seq cohorts with our single-cell data. TCGA COAD and READ cohort masked somatic mutation data and gene expression data was obtained from the GDC portal, survival data from Liu *et al*., 2018^121^. AC-ICAM mutation, survival, and gene expression data was obtained from the paper supplements. For mutations Shears was run with the covariates dataset (AC-ICAM/TCGA) and type (COAD/READ). Shears for survival data was run with the covariates tumor_stage (early/late), age_scaled, and dataset.

### Quantification and statistical analysis

Statistical analysis was performed using the statsmodels library in Python (scRNA-seq data) or GraphPad Prism (flow cytometry and imaging data) using a linear model, t-test or wilcoxon test as appropriate. Single cell-data were aggregated into pseudobulk samples by biological replicates. Compositional analysis of cell-type fractions was performed using scCODA; survival analysis using the Kaplan-Meier formula in R. P-values for untargeted analysis (DE genes, TFs, or pathways) were FDR-adjusted. Significance levels and more details on the statistical tests are indicated in the figure captions.

### Flow Cytometry staining of human samples

Cells isolated from surgically resected CRC tumor and adjacent normal tissues as well as whole blood obtained just prior surgery were stained with a cocktail of the following 20 antibodies (CD3, CD8, CD4, CD14, CD16, CD19, CD15, CD28, CD34, CD31, CD38, CD45, CD56, CD90, CD123, CD161, CD193, CD326, HLA-DR, TCRgd) to define all cell populations of interest. After washing and addition of 5 µL 7-AAD, cells were measured on a FACSymphony A5 flow cytometer (BD Biosciences). All source data of the aforementioned antibodies is provided in the key resources table, applied fluorochromes are listed in Suppl. Table S2. Data were analyzed using FlowJo v10.7 software. For details of the gating strategy, see Figure S8.

### Multiplex immunofluorescence of human samples

CRC tumor and tumor-adjacent tissue samples were fixed in 4% formalin and embedded in paraffin. Three-micrometer sections were used for immunofluorescence staining. The immunofluorescence staining on formalin-fixed paraffin-embedded (FFPE) tissue was conducted using the Opal 6-Plex Detection Kit (Akoya Biosciences). A multiplex panel of immune markers was established utilizing antibodies against CD15 (1:200, clone MMA, Abcam), LOX-1/OLR1 (1:200, polyclonal, Sigma-Aldrich), and cytokeratin (1:1000, clone AE1/AE3, Dako; clone C-11, Abcam). The staining procedure was performed using an automated staining system (BOND-RX, Leica Biosystems). Markers were applied sequentially and paired with their respective Opal fluorophores (CD15 – Opal650, LOX-1/OLR1 – Opal570, cytokeratin – Opal690). To visualize cell nuclei, the tissue was stained with 4’,6-diamidino-2-phenylindole (spectral DAPI, Akoya Biosciences). The stained slides were scanned using the Mantra 2 Quantitative Pathology Workstation (Akoya Biosciences), and representative images were acquired using the Mantra Snap software version 1.0.4. Spectral unmixing, multispectral image analysis, and cell phenotyping were carried out using the inForm Tissue Analysis Software version 2.4.10 (Akoya Biosciences). Pseudo-DAB staining projections were achieved by converting fluorescent channels to pseudo-brightfield images using InForm, where fluorescent markers are displayed as pseudo-DAB staining and DAPI as hematoxylin.

### Sample preparation for Xenium in situ workflow

Sections were prepared according to manufacturer’s instructions (Tissue Preparation Guide Demonstrated Protocol CG000578) at the Department of Anatomy, Histology and Embryology, Medical University of Innsbruck. 5 µm sections were sectioned from FFPE tissue blocks with a microtome (Epredia, HM355S), floated in a 45°C water bath, and adhered to Xenium slides (10x Genomics). Samples were baked at 42°C for 3 hours and stored in a desiccator at room temperature (20°C ± 2°C), overnight, followed by deparaffinization and decrosslinking, according to manufacturer’s instructions (Deparaffinization & Decrosslinking Demonstrated Protocol CG000580).

### Xenium in situ workflow and data acquisition

Xenium samples were processed according to manufacturer’s protocols (Xenium In Situ Gene Expression with Morphology-based Cell Segmentation Staining User Guide CG0000749 and Decoding & Imaging, User Guide CG000584). Samples were stained utilizing the predesigned Human Immuno-Oncology panel from 10x genomics (PN 1000654), as they became available from the manufacturer. Slides were stained with a Xenium Cell Segmentation kit according to manufacturer’s instructions (User Guide CG0000749). Briefly, probes of the immuno-oncology panel were hybridized overnight before rolling circle amplification and native protein autofluorescence was reduced with a chemical autofluorescence quencher. Slides were processed on a 10x Xenium Analyzer, with ROIs selected to cover the selected regions. The Xenium Onboard Analysis pipeline v2.0.0 (10x Genomics) was run directly on the instrument for imaging processing, cell segmentation, image registration, decoding, deduplication, and secondary analysis, as previously described^122^.

### H&E staining of human samples

After processing tissue sections through the Xenium workflow, the same sections on the Xenium slide were used for H&E staining, according to the manufacturer’s instructions (Post-Xenium Analyzer H&E Staining CG000613) and imaged using a PANNORAMIC slide scanners (Institute for Pathology, Neuropathology & Molecular Pathology, Medical University of Innsbruck).

### Immunohistochemistry of human samples

Consecutive FFPE tissue slices of CRC tumor core, invasive margin and adjacent normal samples (n = 11) were stained with the following antibody for immunohistochemistry (IHC): CONFIRM anti-CD15 (clone MMA, Mouse Monoclonal, Primary Antibody, ready-to-use (RUI), IVD (Ventana, Roche). Staining was made on a BenchMark Ultra instrument (Ventana, Roche). The whole slides were scanned with the VENTANA® DP 200 slide scanner (Ventana Roche) and the navify® Digital Pathology platform to retrieve digital slide tissue images. Digital slides were further processed with the QuPath software v0.5.1 and ImageJ software v1.54g.

Post Xenium and H&E staining, CRC tumor core and invasive margin samples (n = 5) were stained with CD15 antibody and assessed by IHC on an automated staining device (DAKO Omnis, Agilent) by using a ready-to-use kit (mouse anti-human, clone Carb-3, Agilent). Normal human tonsil and normal skeletal muscle were included as positive and negative controls in each staining run.

### Xenium in situ data analysis

Sample areas were manually selected on the Xenium Analyzer or in the Xenium Explorer software. Xenium output bundles were further processed with scanpy^105^, squidpy^123^ and the SpatialData framework^124^. Samples from different slides were merged and spatially aligned. Cells were filtered to have a minimum number of 15 transcripts, less than two control codewords, and at least four spatial neighbors within 50 μm, resulting in a dataset of 3.7 million cells. Integration of samples was done on the filtered counts matrix with an scvi model of one hidden layer and 30 latent components. Leiden clustering was performed on the integrated latent space. To annotate leiden clusters, we used a three-step approach: First, we utilized tangram^125^ in cluster mode to predict cell type labels with both a downsampled version of the complete CRC atlas and with only the MUI cohort, which consists of single-cell RNA sequencing samples from matched patients. scSampler^126^ was used for downsampling of the complete atlas. To generate a consensus annotation, we mapped the middle and fine cluster annotations from the references to single cells of the spatial atlas, using the batch-corrected counts as expression profiles. Leiden clusters at resolution 2.5 were then annotated by the most abundant predicted middle cell type label and at resolution 3.8 by the most abundant predicted fine cell type label. Second, we used a Wilcoxon test to calculate the top markers per cluster to identify cell types not present in the reference data (i.e., smooth muscle cells). Third, we checked the consistency of the annotated cell types with their spatial localization. To identify cell niches, NicheCompass^127^ was run on the raw counts with the ‘gcnconv’ encoder and each sample as batch factor. OmniPath^128^ and NicheNet^129^ were used as prior gene programs. The input spatial neighbors graph was calculated with squidpy using Delaunay triangulation for cell centroids within a radius of 50 µm. Leiden clustering was then performed at different resolutions on the niches latent space to identify distinct spatial niches. Marker genes for each cell type within a given niche were obtained using a Wilcoxon test with pre-test filtering to remove spillover transcripts from adjacent cells. Neutrophil clusters were identified from the spatial neighbors graph by first defining cluster seeds as cells of cell type ‘Neutrophil’ with at least three homotypic neighbors. Cluster seeds were subsequently expanded in ten iterations to add all neutrophils adjacent to seed cells to the respective cluster. To investigate cell-cell-communication pathways within cell niches, the steady-state version of LIANA+^33^ was run for cell types with at least 100 cells in a given niche, consensus resources, default methods and 10,000 permutations.

### Tissue microarrays and IF image acquisition

Five tissue microarrays (TMAs) containing a total of 549 valid cores were processed and stained for HLA-DR and CD15 using immunofluorescence (IF). In brief, slides have been deparaffinized, processed by heat-induced epitope retrieval and incubated with anti-HLA-DR (1:250, Abcam) and anti-CD15 (Dako/Agilent). Staining has been visualized by fluorochrome-labeled secondary antibodies (ThermoFisher); for HLA-DR staining, HLA-DR antibody has been pre-labeled using FlexAble CoraLite Plus 555 (Proteintech) according to the manufacturer’s instructions. Images of the stained TMAs and whole slides were acquired in .czi format using AxioScan Z.1 (Zeiss).

### IF image preprocessing and core extraction

The .czi images were converted to a compatible format (.tif) for downstream analysis using Fiji (ImageJ, v1.54f). Images were carefully inspected during this step to ensure no loss of resolution in any of the RGB layers. Individual TMA cores were extracted from the pre-processed images using QuPath (v0.5.1), an open-source software platform for digital pathology. The cores were manually annotated to exclude damaged or improperly stained regions, and only valid cores were included in subsequent analyses.

### Cell identification and quantification in IF images

A customized image analysis pipeline was developed in CellProfiler (v4.2.8) to identify and quantify nuclei and marker-positive cells for CD15 and HLA-DR. The pipeline consisted of the following steps:

1. Nuclear Identification: Nuclei were segmented and identified based on fluorescence intensity.
2. Marker Co-Localization: Positive staining for CD15 and HLA-DR was identified and quantified, with nuclei serving as reference points for spatial localization and co-expression analysis.

Output: The pipeline produced quantitative data on nuclei counts, their spatial position and the number of nuclei positive for each marker, based on staining intensity to distinguish specific marker expression from background fluorescence. Quantitative data, including total nuclei count, all CD15-positive cells, all HLA-DR-positive cells and co-expression counts, were exported from CellProfiler for statistical analysis.

### Imaging mass cytometry and data analysis

Imaging mass cytometry was performed on tissue microarrays from microsatellite-stable CRC patients as previously described^35^, using the antibody panel provided in Suppl. Table S2. Images were obtained from a total of 37 patient samples, including intratumoral regions from 27 patients and tumor margin regions from 29 patients. Among these, both intratumoral and tumor margin regions were imaged for 19 patients. Per sample, one or two 1000x1000 µm regions of interest were ablated, depending on tissue availability. Images were visually inspected and exported to ome.tiff using MCD viewer (standard biotools). Image normalization, cell segmentation and phenotype identification were performed as described previously^35^ and steps were validated throughout on the original images.

### Spatial analysis of neutrophil niches for IF and IMC data

To analyze the spatial organization in tumor and margin regions, we constructed Voronoi diagrams based on the center coordinates of all detected nuclei. Each Voronoi region was then assigned the corresponding cell type. The DBscan clustering algorithm was used to find niches consisting of at least 3 neutrophils (granulocytes) in a maximum pairwise distance of 15 µm. For these niches all other cell types that share a Voronoi ridge (edge) with a neutrophil were searched and added as “niche members”. To determine if the number of detected niches differed significantly from what would be expected by chance, a Monte Carlo simulation with 1,000 iterations was conducted. In this simulation, the cell type labels for each cell on the imaged slide was randomly assigned while maintaining the number of cells for each type constant and preserving the overall spatial arrangement. A z-score and p-value were then calculated (see Suppl. Table S4).

### Survival analysis

Survival analysis was performed to assess the impact of the number of neutrophil niches on overall survival. Patients were stratified into “high” and “low” groups based on an optimal cutoff that minimizes the p-value or by or by setting it to 0.66. Kaplan-Meier survival curves were generated for each group using the KaplanMeierFitter function from the lifelines Python package. The log-rank test, implemented via the logrank_test function from the same package, was used to compare survival distributions between the two groups. Cox proportional hazards regression was employed using the CoxPHFitter function (lifelines) to calculate hazard ratios (HR) and 95% confidence intervals (CI).

### Mice

C57BL/6J (B6J, stock no. 000664) mice were obtained from The Jackson laboratory and used for injections of AKPS organoids, or kept uninjected as non-tumour bearing control mice. All mouse experiments were performed under the approval of Cantonal Veterinary Office of Zurich, Switzerland and in accordance with Swiss guidelines. All mice were maintained on a 12h light and 12h dark schedule, and water and chow were provided ad libitum.

### Colonoscopy submucosal injection of AKPS organoids

*Apc*-LOF, *p53*-LOF, *Kras*-GOF, *Smad4*-LOF (AKPS) tumor organoids were generated from AKP organoids harbouring a RNP-mediated Smad4 mutation introduced by CRISPR-Cas9^36^. Organoids were cultured in Matrigel and passaged every 2-3 days^130^. Colonoscopy-guided orthotopic cell transplantation was done as described previously with modifications^131^. Briefly, following mechanical dissociation, the equivalent of one Matrigel done was resuspended in 50 mL OptiMEM (Gibco) to equate to one injection in one mouse. 8-to 10-week-old males were anesthetized with isoflurane and placed facing up on a 37°C heating pad. Fecal content was flushed from the colon with warm DPBS and AKPS organoid solution was injected carefully below the mucosa using a custom needle, syringe and colonoscope. Observation of a visible injection bubble was noted as an indicator for successful injection in the correct location (Figure S6A). Mice were then regularly monitored according to official guidelines until experimental endpoint.

### Tissue dissociation of mouse samples

#### AKPS tumors

Tumors and disseminated tumors from the peritoneal cavity of 5 mice were collected at the study endpoint, pooled respectively and cut into small pieces. Tissue was digested at 37 °C for 50 min in RPMI-1640 containing 15 mM HEPES (H0887 Sigma), 500 U/mL^-1^ type IV (C5138 Sigma) and type VIII collagenase, (C2139 Sigma) and 0.05 mg/mL DNase I (10104159001 Roche). Non-digested tissue was mechanically disrupted with a syringe and cell suspension was pushed through a 70 µm cell strainer. Flow-through was topped with DPBS buffer containing 2% BSA, 100 U mL/^-1^ penicillin/streptomycin (P0781 Sigma) and 5 mM EDTA and cells were spun prior and resuspended prior analysis.

#### Blood from naïve and AKPS tumor-bearing mice

Blood was collected from mice post-mortem by cardiac puncture in 2% BSA 5 mM EDTA, followed by red blood cell lysis with distilled cold water for 30 sec. Blood from 3 naïve and 5 AKPS tumor-bearing mice were pooled respectively.

#### Colon from naive mice and adjacent colon to AKPS tumor

Colonic tissue from 5 mice per condition was collected, cleaned and opened longitudinally, prior to washing with PBS. Tissues were cut into small pieces, pooled per condition and washed twice while shaking for 25 min at 37 °C with HBSS 2% BSA, 100 U mL/^-1^ penicillin/streptomycin (P0781 Sigma) and 5 mM EDTA. Tissue was digested at 37 °C for 50 min in RPMI-1640 containing 15 mM HEPES (H0887 Sigma), 500 U/mL^-1^ type IV (C5138 Sigma) and type VIII collagenase, (C2139 Sigma) and 0.05 mg/mL DNase I (10104159001 Roche). Non-digested tissue was mechanically disrupted with a syringe and cell suspension was pushed through a 70 µm cell strainer. Cells were centrifuged for 8 min and layered onto a 40/80% Percoll (17089101 Cytiva) gradient (18 min, 2.100 *g*, 20 °C, no brake). The interphase was collected and washed in DPBS.

#### Bone marrow from naïve and AKPS tumor-bearing mice

Tibia and femur were collected and flushed with RPMI medium using a 23-gauge needle. The content was filtered through a 40 µm strainer followed by red blood cell lysis via incubation in ice-cold distilled water for 30 sec. Bone marrow from 3 naïve and 5 AKPS tumor-bearing mice were pooled respectively.

### Flow cytometry staining of mouse samples

#### Staining

Cells were surface stained in DPBS at 4 °C for 30 minutes with fixable viability dye eFluor 780, and combinations of the following listed antibodies (1:200, all from BioLegend; unless otherwise stated): CD11b BV510 (M1/70, 101263), CD45 BV650 (30-F11,103151), CD80 BV605 (1:100, 16-10A1, 104729), Ly6G Percp-Cy5.5 (1A8, 127616), MHCII AF700 (M5/114.15.2,107622), PD-L1 PE-Cy7 (1:100, 10F.9G2, 124314), Siglec-F BV421 (E50-2440, 552681 BD Biosciences). Samples were acquired on a LSRII Fortessa (BD Biosciences).

#### Data analysis

Analysis of flow cytometry data was performed using FlowJo Software. Relative cell frequencies, cell counts, or Mean Fluorescent Intensities (MFIs) were subsequently plotted in graphical formats using GraphPad Prism.

#### Statistical analysis

Statistical analyses were performed using GraphPad Prism. For statistical comparisons of two groups, two-tailed unpaired t-tests were used. For comparisons of more than two groups, a one-way analysis of variance (ANOVA) was used. Differences were considered statistically significant when P < 0.05.

### Single-cell RNA sequencing of mouse neutrophils - experimental

#### CD45+ cell enrichment using magnetic beads

CD45 positive cells were enriched using anti-CD45 microbeads (130-052-301 Miltenyi Biotec) according to the manufacturer’s instructions.

#### Cell hashing

Samples were pooled for shared scRNAseq runs after labeling with sample tags (633793 BD Mouse Single-Cell Multiplexing Kit). Thereby, 10^6^ cells from each sample were incubated in a staining buffer (DPBS, 1% BSA, 1% EDTA) and the respective sample tag for 20 min at room temperature (20°C ± 2°C). After incubation, cells were washed twice with staining buffer before counting. From each sample 10-20.000 cells were pooled to have a total of around 60-80.000 cells for single cell capture.

#### Single cell capture, library preparation and sequencing

Single cell capture was done using the BD Rhapsody Express Single-Cell Analysis System (BD Biosciences) following the manufacturer’s protocol. In short, cells were resuspended in 650 µL BD Sample Buffer supplemented with Rnase inhibitors (1:1.000 SUPERase 20 U µL^-1^, AM2694 Thermo Fisher Scientific; NxGen Rnase Inhibitor 40 U µL^-1^, 30281-2 Lucigen) and loaded on a cartridge. To extract their RNA capture beads (v1 for tumor, disseminated and adjacent colon; enhanced beads for colon, blood and bone marrow) were added, followed by cell lysis. After single cell capture, libraries from the cDNA and the sample tags were prepared using the BD Rhapsody Whole Transcriptome Analysis Amplification Kit (633801 BD Biosciences). Libraries were then quantified using a Qubit Fluorometer (dsDNA HS kit: Q32851 Thermo Fisher Scientific) and a TapeStation 4200 (HS D500 types and reagents, 5067-5592 Agilent Technologies). Sequencing was performed on a NovaSeq6000 system (Illumina) with SP100 Reagents (100 cycles) and PhiX spike-in (20% for v1, 1% for enhanced beads) with a depth of around 20.000 reads/cell for WTA and 500-1000 reads/cell for the sample tags.

### Single-cell RNA sequencing of mouse neutrophils - data analysis

#### Pre-processing, normalization, clustering and data integration

Demultiplexing was done using Bcl2fastq (v2.20.0433, Illumina). FASTQ files were then further processed using the BD Rhapsody^TM^ WTA analysis pipeline (v1.12.1) on the Seven Bridge Genomics platform with default settings. Gene mapping was done with a STAR index generated using STAR^132^ (v2.5.2b) with the gene code GRCm38 v25. R (v4.3) and Python (v3) were used for the downstream analysis. If not specified, the Seurat^133^ (v5.1.0) R package was used. After the first quality assessment, cells with fewer than 200 gene features, more than 5000 gene features and more than 25% of mitochondrial genes were removed from the dataset. Additionally, the R package decontX^134^ (v1.0.0) was applied to adjust the dataset for cell free RNA. This approach uses a Bayesian model to estimate and remove cell free RNA contaminations. Data was log normalized and scaled with the Seurat functions ‘NormalizeData’ and ‘ScaleData’. Thereby, the number of features, UMI counts and percentage of mitochondrial genes were regressed out. The top 2.000 variable feature genes were used for principal component analysis (PCA) followed by clustering using the Louvain algorithm with multi level refinements. The colon and tumor samples were processed separate from the blood and bone marrow samples. To account for batch effects in the colon and tumor samples the fast mutual nearest neighbors (FastMNN) algorithm (Seurat function ‘FastMNNIntegration’) was applied. The blood and bone marrow samples were merged without batch effect correction.

#### Annotation, extraction and integration of neutrophils

Neutrophil clusters were identified using on the one hand neutrophil specific marker genes (*S100a8, S100a9*) and an automatic annotation approach called SingleR^135^ (v2.4.1) with the murine ImmGen^136^ (Immunological Genome Project) as reference which was loaded with the celldex^135^ (v1.12.0) R package. Neutrophil clusters were then extracted and integrated from all tissues using the FastMNN approach mentioned before.

#### Analysis of human neutrophil markers in mouse data

To analyze the expression of marker genes identified in human neutrophil clusters, the genes were first converted to mouse gene symbols using the orthogene Bioconductor package (v1.8.0) and then plotted in a heatmap using pheatmap (v1.0.12).

#### Pseudotime trajectory analysis with Slingshot

A pseudotime trajectory was inferred on the mnn.clusters of the combined mouse neutrophils dataset (bone marrow, blood and colon) using the slingshot^137^ (v2.10.0) R package. Therefore, the Seurat object was first converted to a Single Cell Expression (SCE) object using sceasy R package (v0.0.7), which was then used for the slingshot analysis on the UMAP dimensional space. In brief, by minimum spanning tree (MST) construction using the ‘getLineages’ function, a global lineage structure was identified and pseudotime variables were inferred using the ‘getCurves’ function. To compare the pseudotime between tissue, the cellular distribution was visualized in a density plot across pseudotime. Transcriptional changes along the trajectory were analyzed using tradeSeq^138^ (v1.16.0). Thereby, a negative binomial generalized additive model (NB-GAM) was applied to smooth the gene expression of the top 2000 variable features (Seurat function ‘VariableFeatures’) into the pseudotime lineage. This was done with the tradeSeq function ‘fitGAM’ for 5 knots. The knots represent points where smoothers are joined together. To statistically test if a certain gene is associated with the lineage the ‘associationTest’ function was applied which is based on a Wald test and p-values were additionally corrected for multiple comparisons using the false discovery rate (FDR) method. Selected genes were then plotted in a heatmap using pheatmap R package (v1.0.12).

#### Differential gene expression analysis

The Seurat function ‘FindMarkers’ with min.pct = 0.25 and logfc.threshold = 0.2 was used on normalized data to analyse DEGs between healthy and tumor bone marrow and blood. Thereby a non-parametric Wilcoxon Rank-Sum test was applied and p-values were corrected for multiple testing using the Bonferroni method.

### Quantification of Neutrophil Activation via Seahorse XF analysis

Human neutrophils were isolated from freshly collected peripheral blood from healthy donors. Neutrophils were purified using the MACSxpress Whole Blood Neutrophil Isolation Kit (Miltenyi Biotec) according to the manufacturer’s instructions. Neutrophils were washed with DPBS (Sigma-Aldrich) and centrifuged at 115 g for 10 minutes. Then, 2 million neutrophils were seeded in a 12-well plate in 1 mL of either RPMI 1640 (Sigma-Aldrich) supplemented with 2 mM L-glutamine (Sigma-Aldrich), 10% FBS (Sigma-Aldrich) and 10 mL/L Penicillin-Streptomycin (Sigma-Aldrich), or with conditioned medium obtained as supernatant of SW900 or CaCo2 cell lines after 48 hours of growth in RPMI (supplemented with L-glutamine, FBS and Penicillin-Streptomycin) to approximately 70% confluency. Neutrophils were cultured for 24 hours at 37°C and 5% CO_2_. Neutrophils were collected, counted, and centrifuged at 450 g for 10 minutes at room temperature (20°C ± 2°C). Cells were resuspended at a concentration of 2.5x10^5^ cells/mL in Seahorse XF RPMI Medium pH 7.4 supplemented with 1 mM pyruvate, 2 mM glutamine, 10 mM glucose (all from Agilent Technologies). Seahorse XFp Cell Culture Miniplates (Agilent Technologies) were pre-coated with poly-L-lysine (Sigma-Aldrich) and 5x10^4^ cells were plated into each well. Miniplates were centrifuged at 300 g for 1 minute with low-brake deceleration. Seahorse XP Neutrophil Activation assay was performed on an XFp instrument with sequential injection of 0.5 µM Rotenone/Antimycin A (Agilent Technologies) and 100 ng/mL phorbol 12-myristate 13-acetate (PMA) (Fisher Scientific) according to the manufacturer’s protocol. Data was processed by Seahorse XFe Wave software (Agilent Technologies) and by GraphPad Prism software (v 10.4; GraphPad Software). Proton efflux rate (PER) is calculated via Wave software as a quantitative measurement of extracellular acidification post run from the real-time acidification data measured during the assay (OCR vs ECAR). Oxygen consumption was calculated from the area under the curve (AUC) of the kinetic range between activation and return to basal rates.

## Key resources table

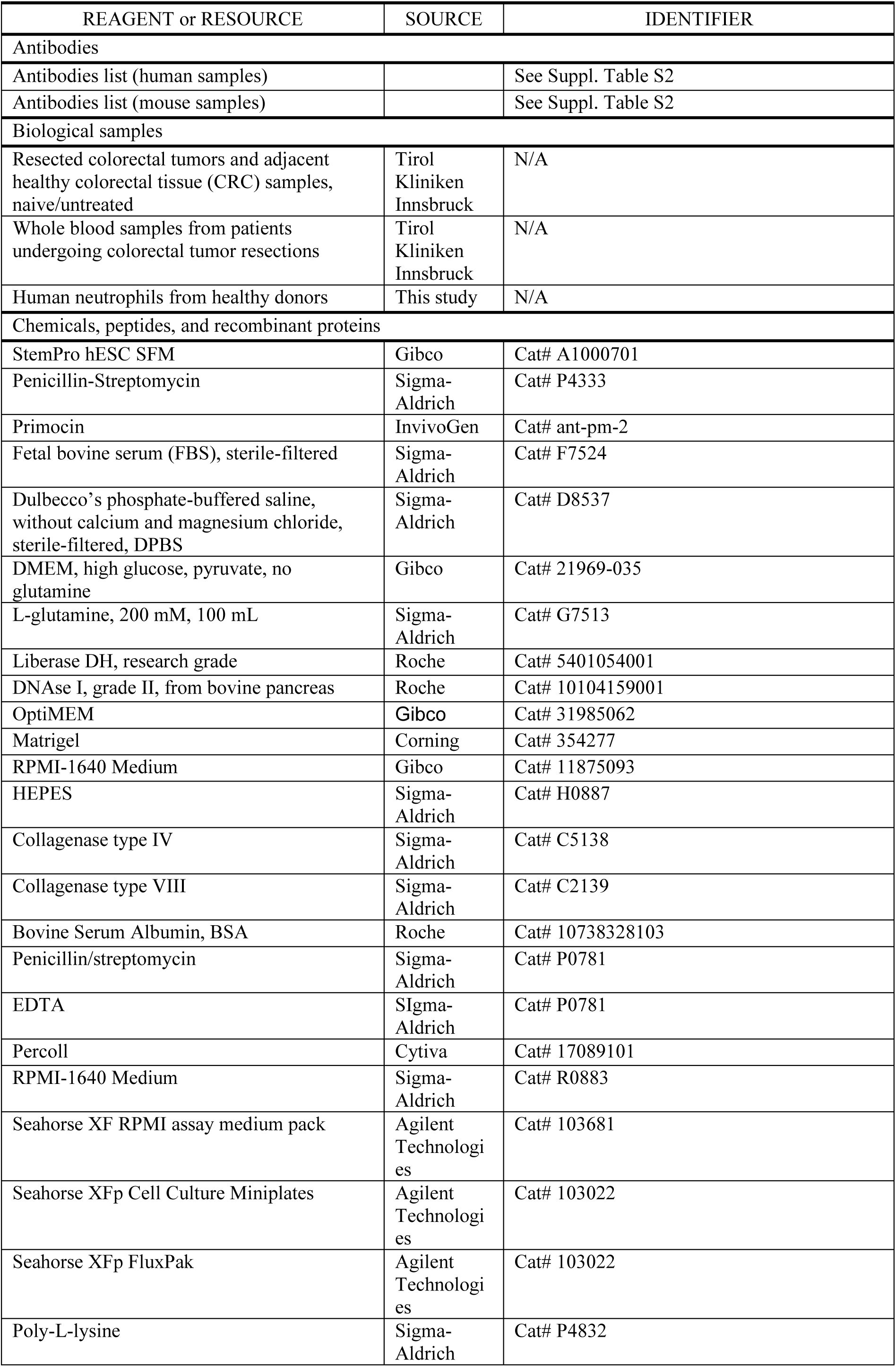
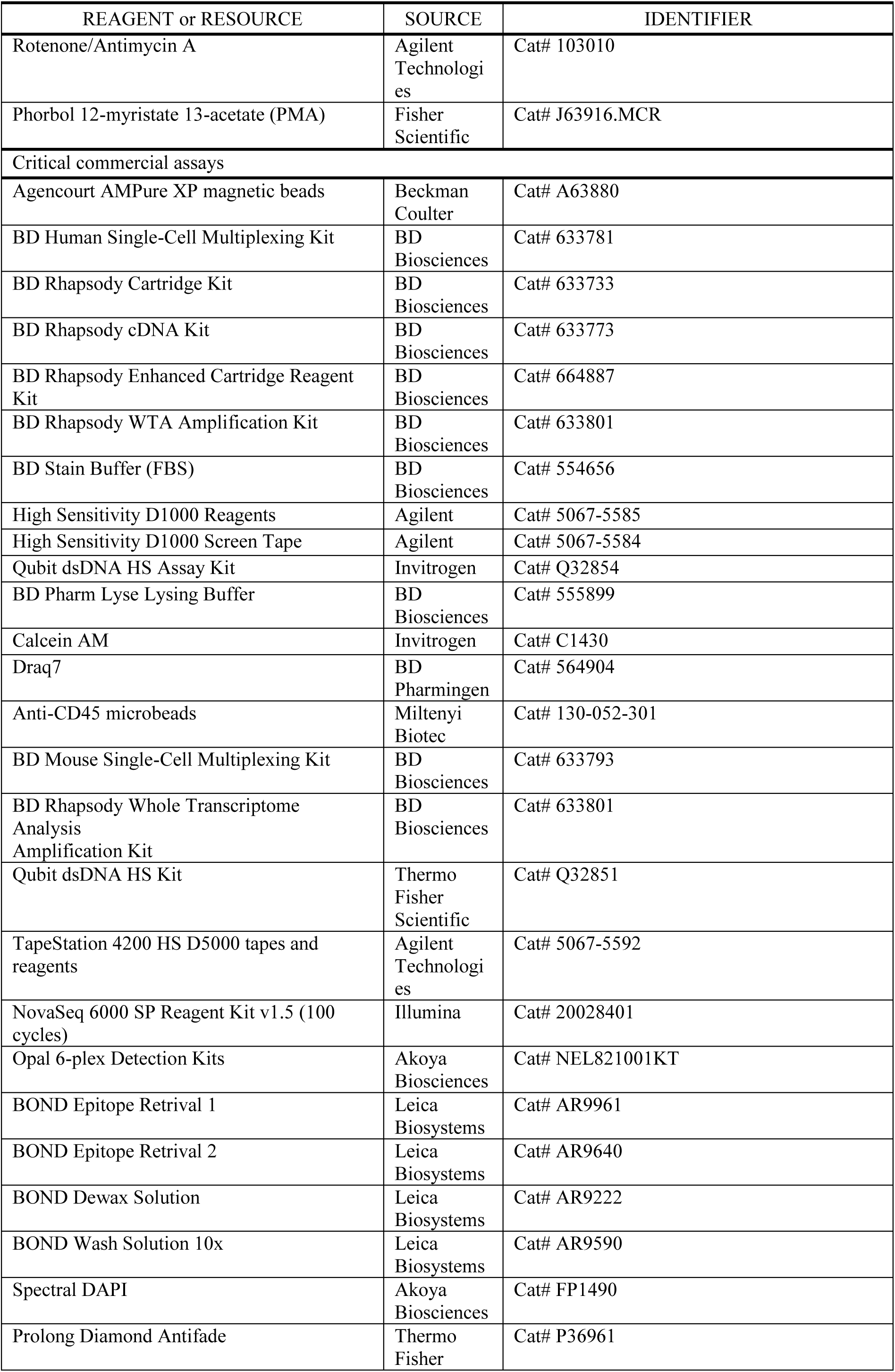
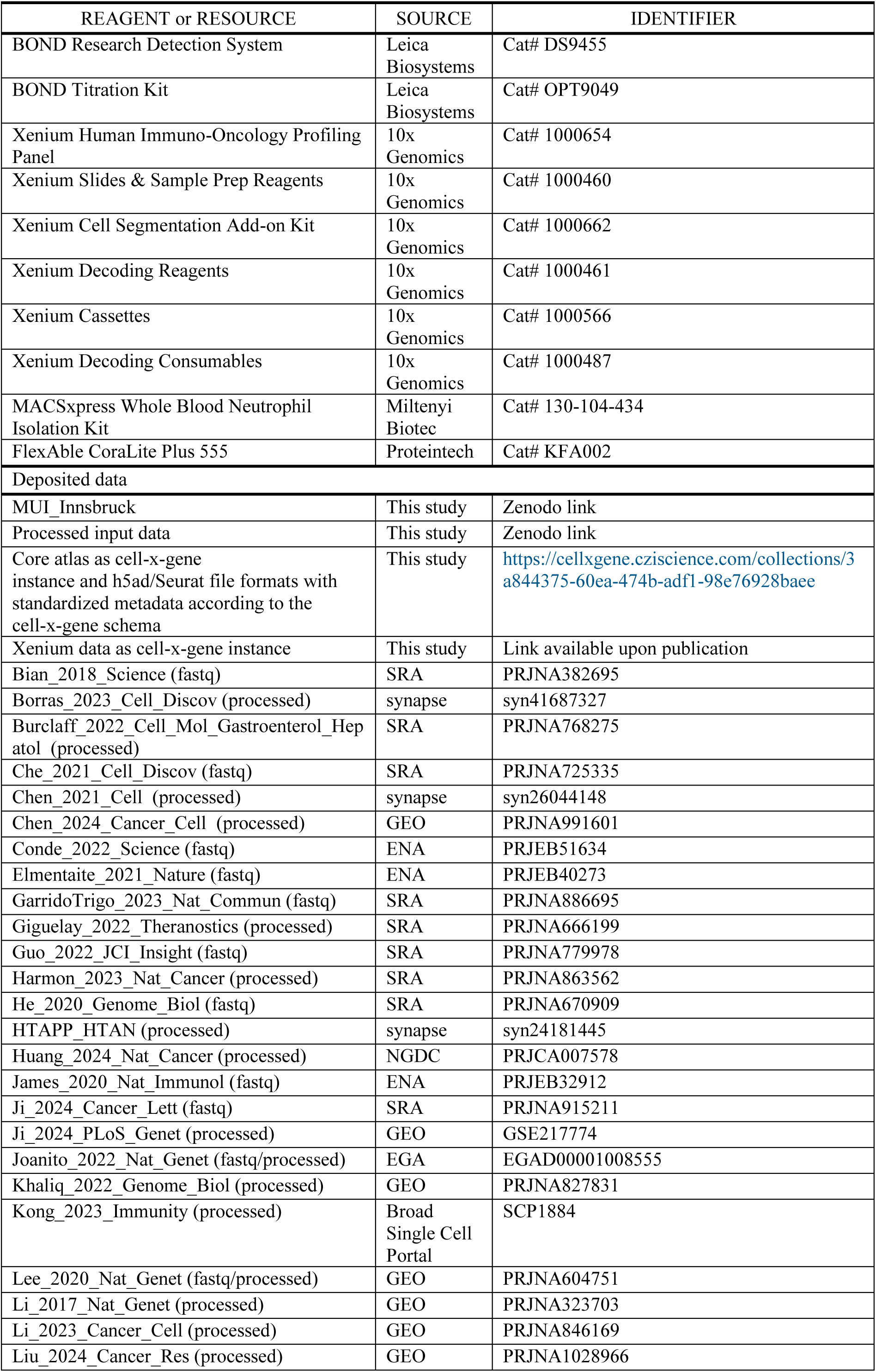
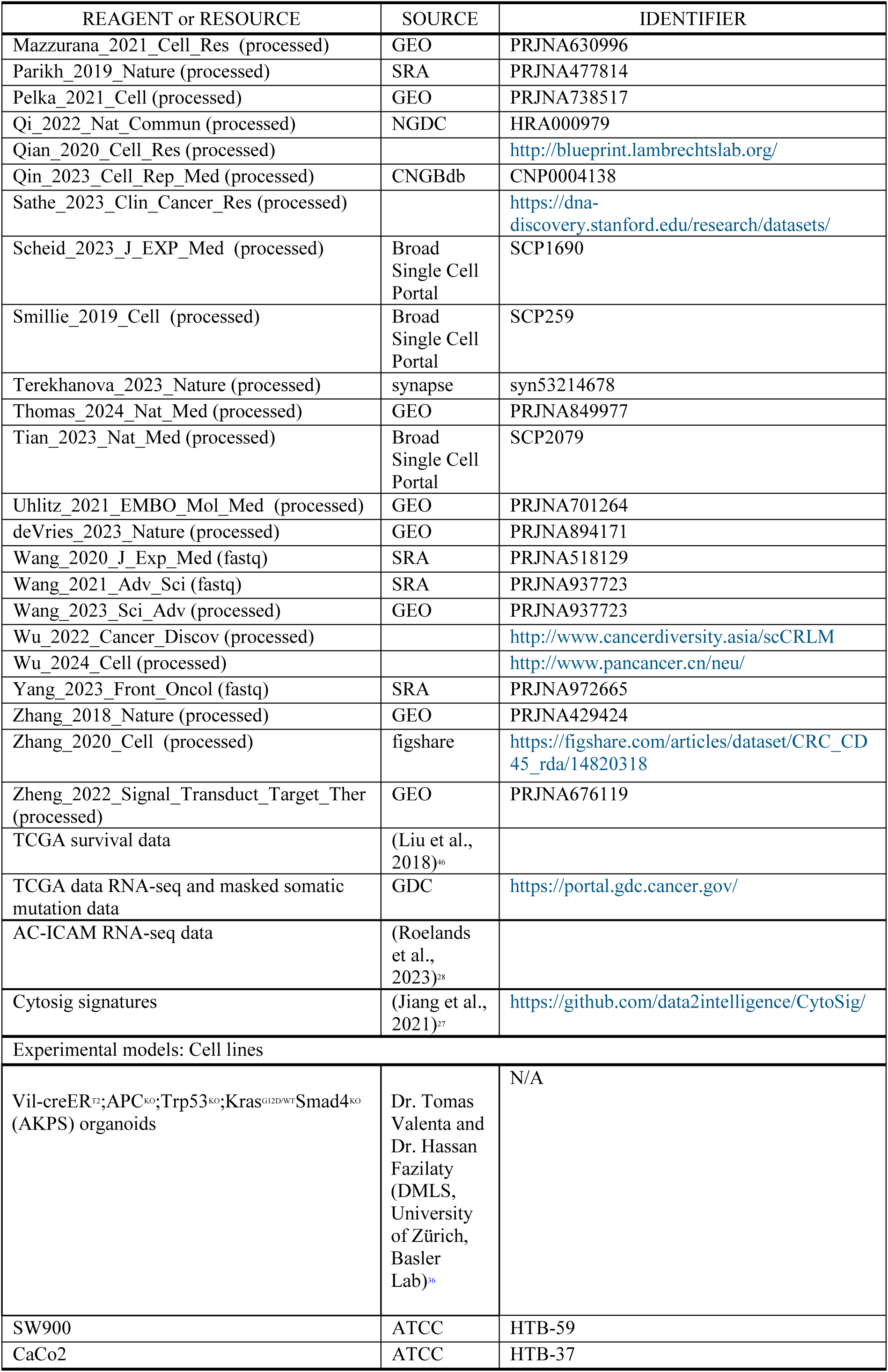
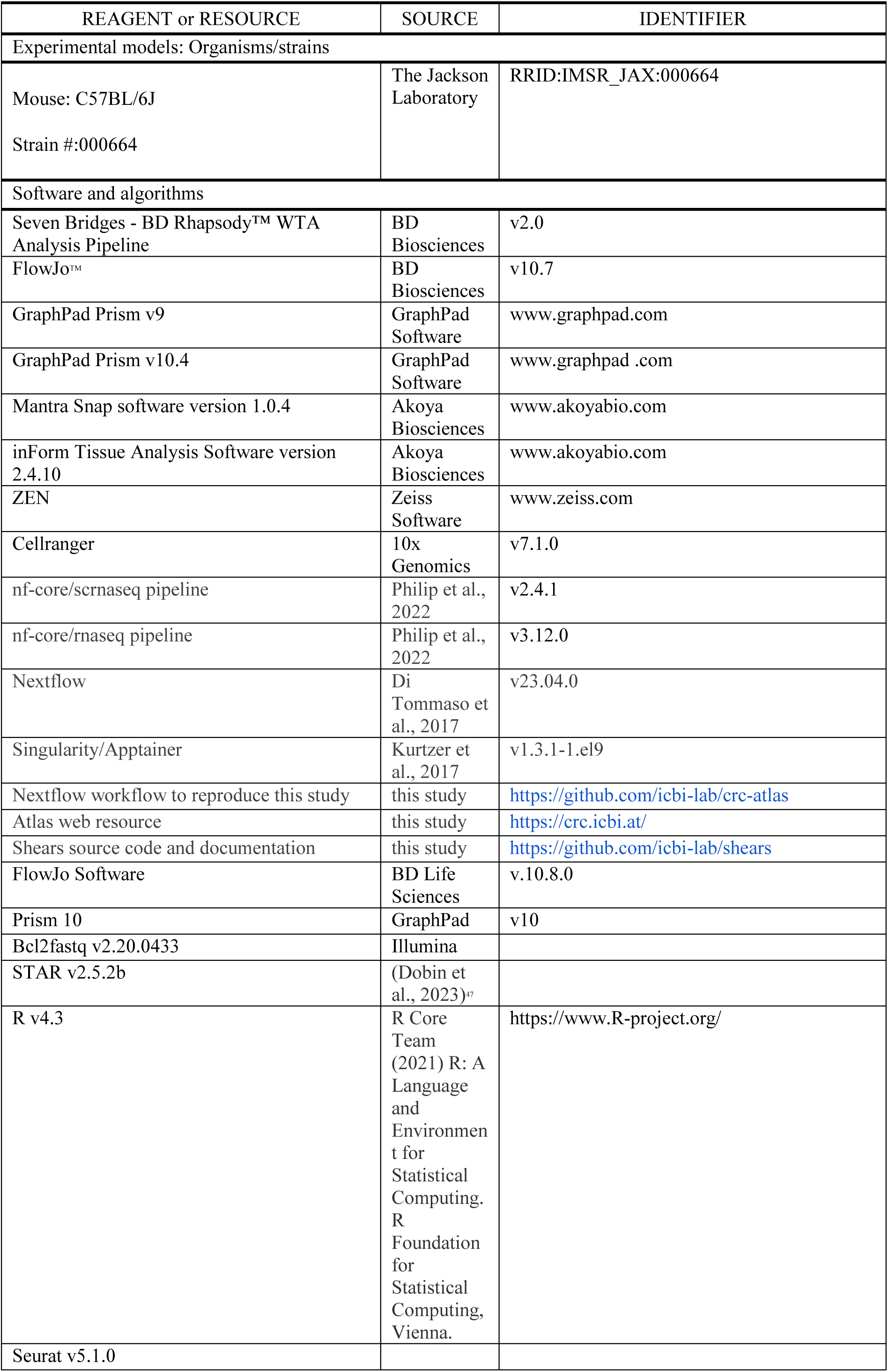
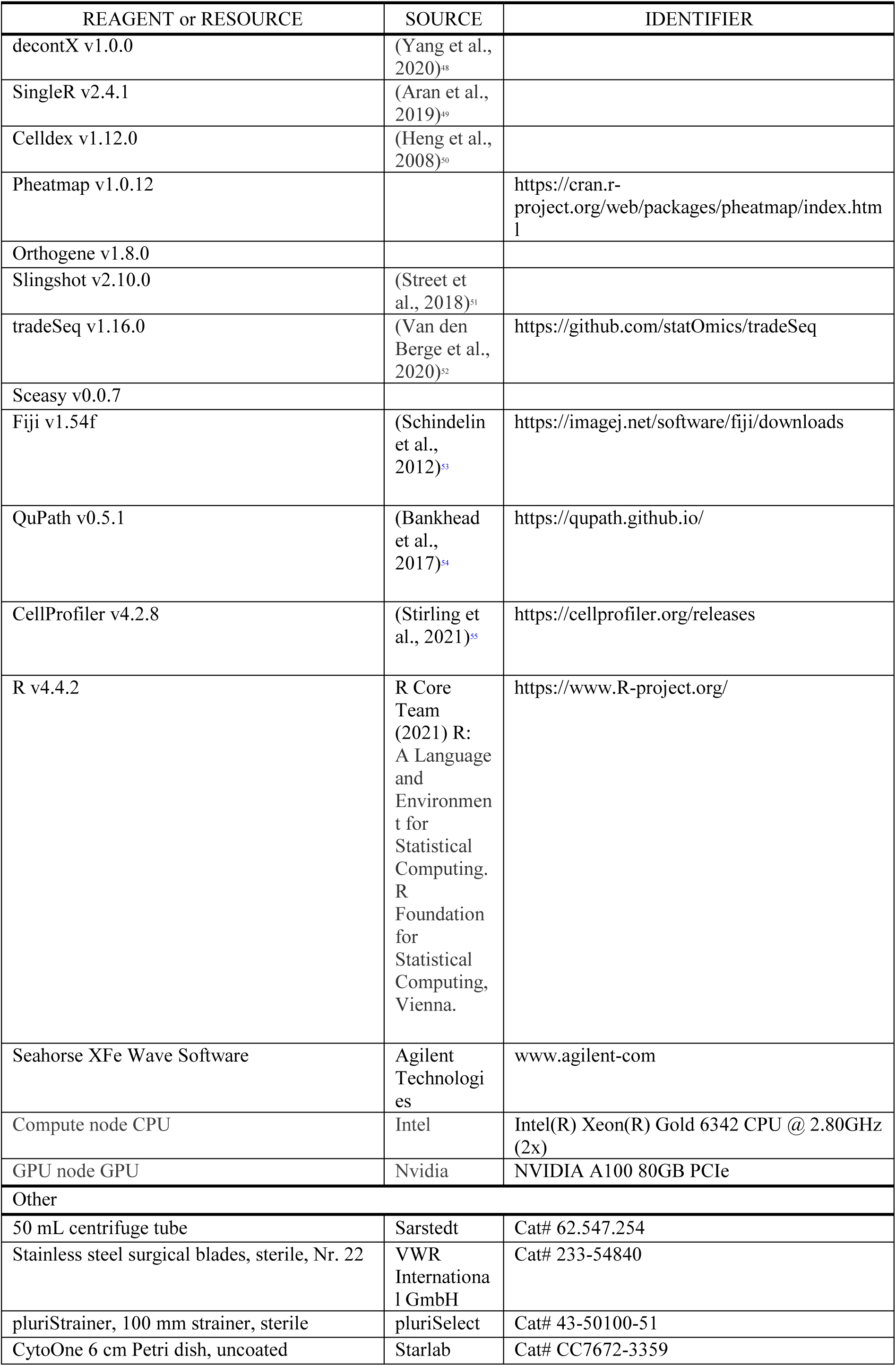
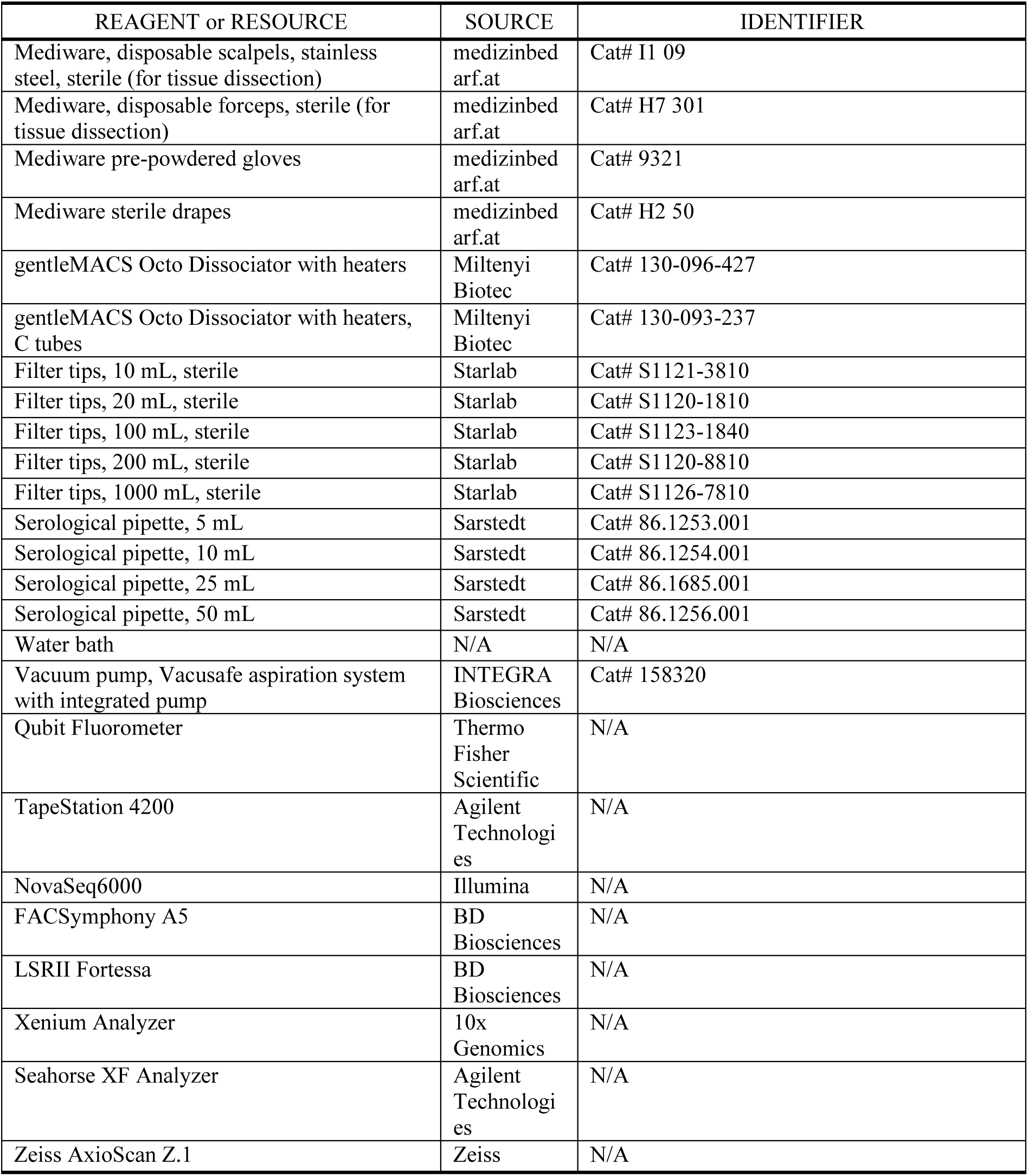

**Figure S1:**
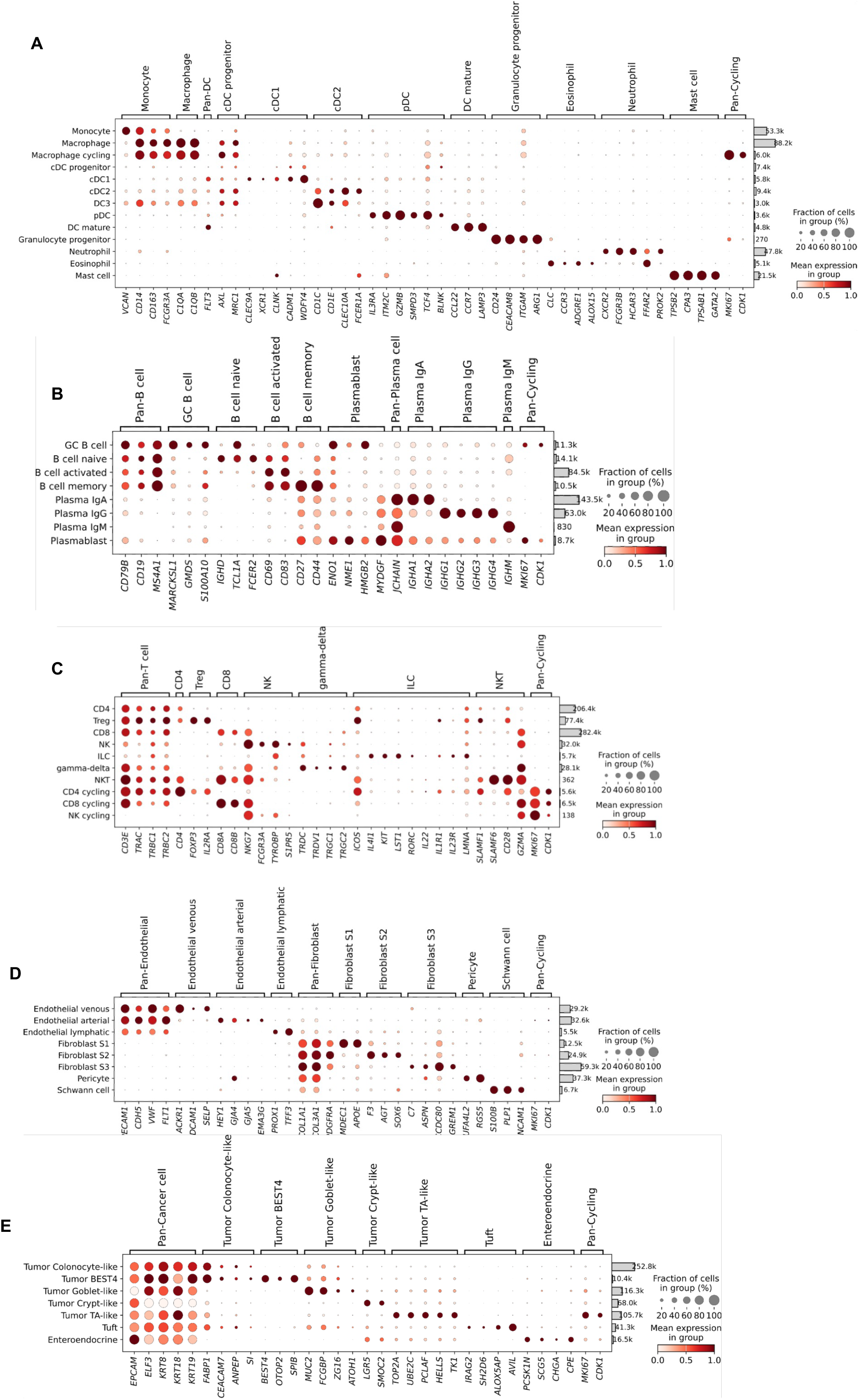
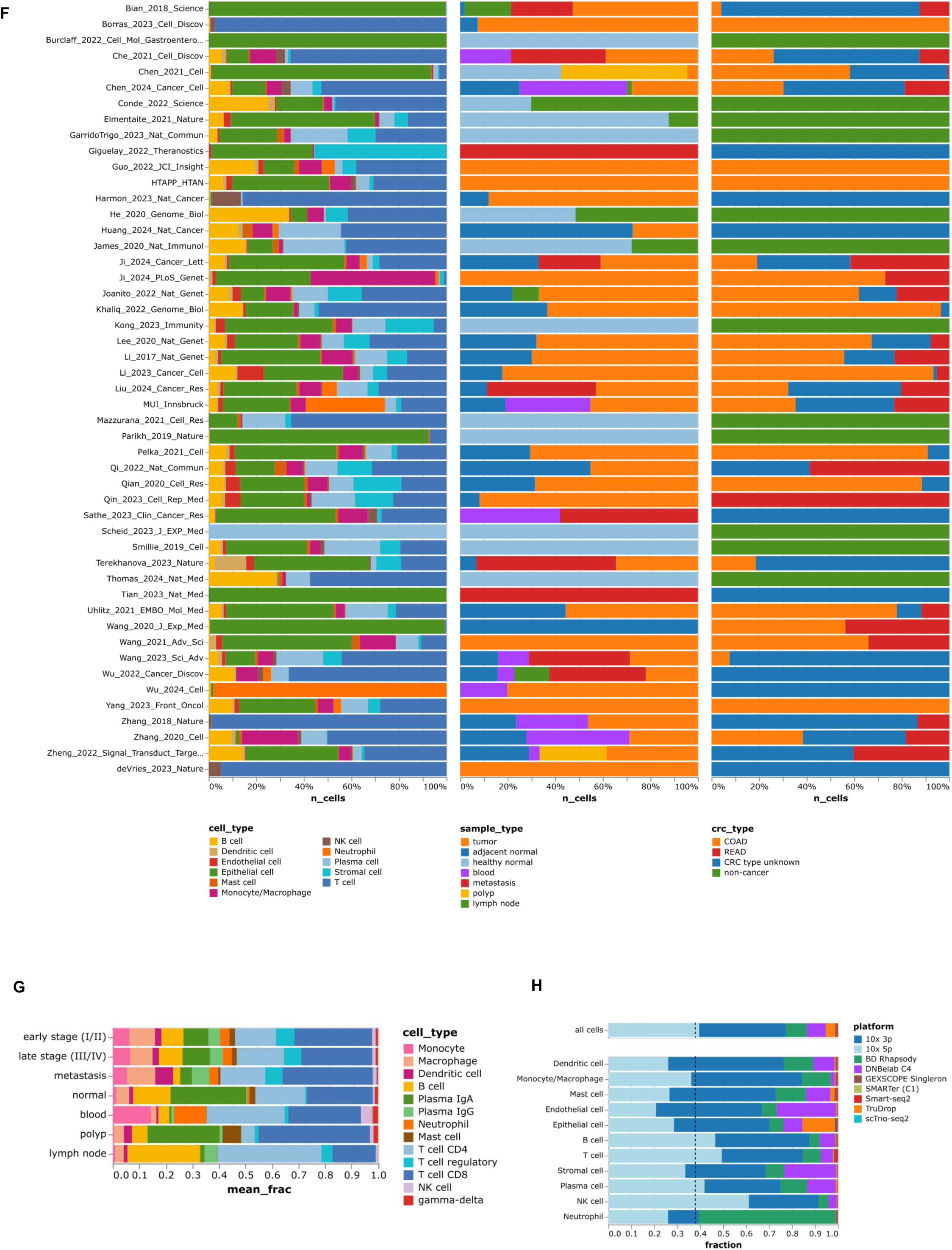
related to Figure 1. Schematic outline of the overall concept of the study including publicly available and generated datasets. Dotplots of cell type marker genes used for cell-type annotation in the (**A**) myeloid lineage **(B)** B/Plasma cell and (**C**) T cell lineages, (**D**) Endothelial and Stromal lineage, and **(E)** Epithelial lineage **(G)** Fraction of cell types, sample types, and CRC types across different datasets, (**H**) Relative cell type proportions of the immune cell subset in the atlas (**I**) Fractions of depicted cell types per scRNAseq platform.

**Figure S2:**
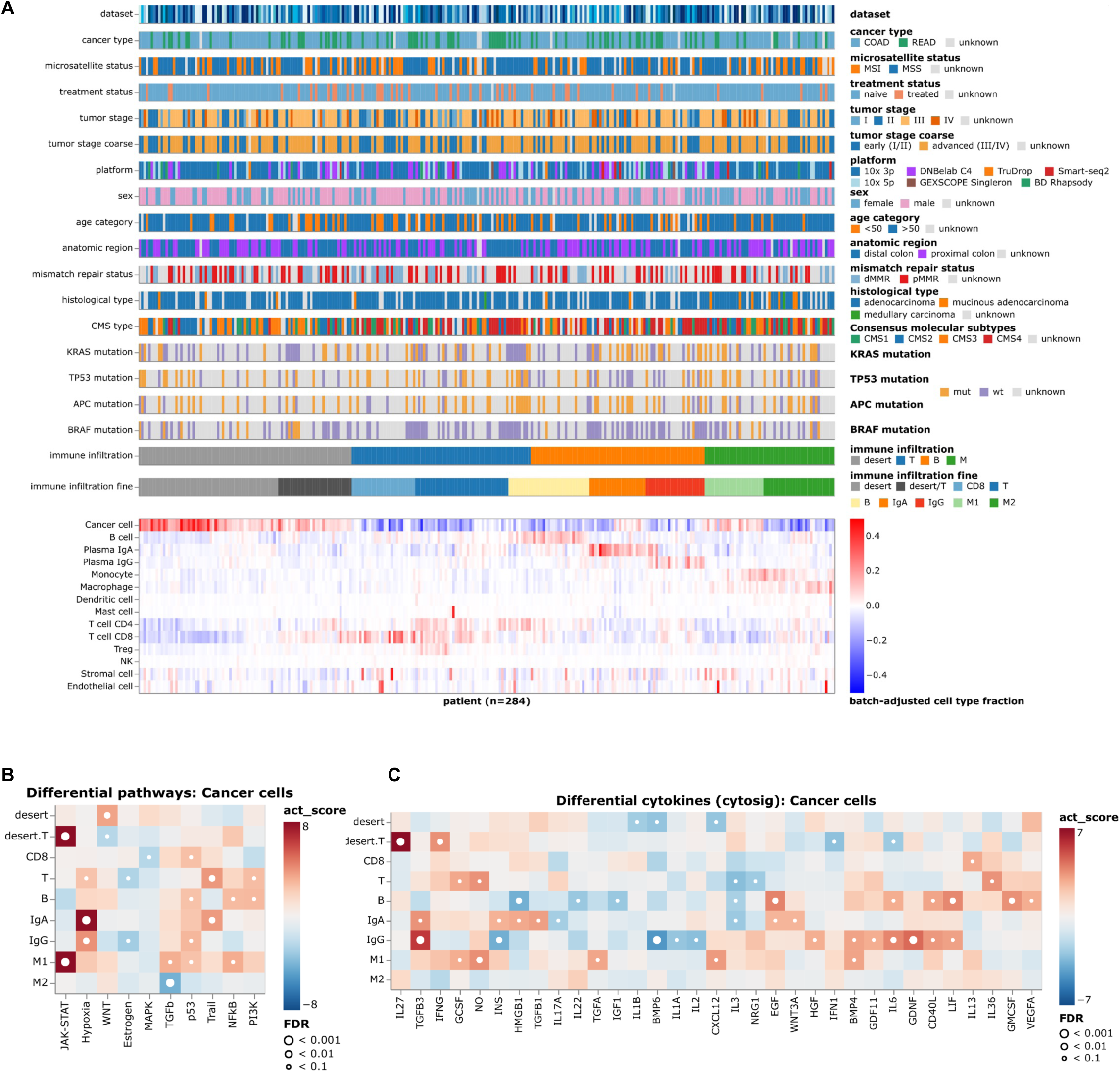
related to Figure 2: CRC patient stratification based on tumor immune phenotypes. (**A**) Immune phenotypes and clinical, genetic features. (**B**) Results of the Progeny analysis. (**C**) Results of the cytosig analysis.

**Figure S3:**
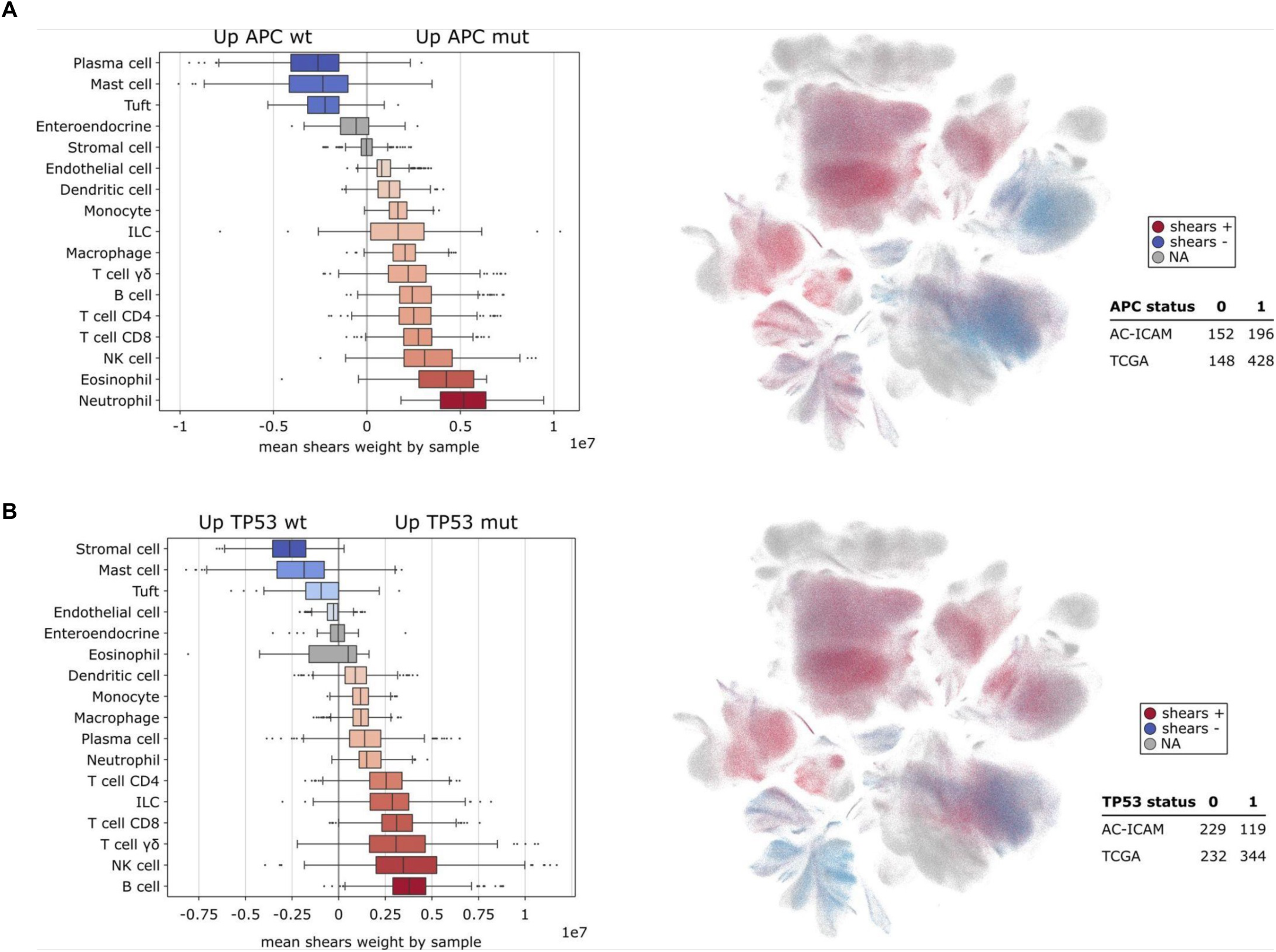
related to Figure 3. Associations of cellular composition, survival, and genotypes. (**A-B**) *SHEARS* analysis showing the association of cellular composition and phenotypic information from bulk RNA-seq data from the combined AC-ICAM and TCGA cohorts. Boxplots summarize the central Shears tendencies for each cell type in relation to the phenotype, with each data point representing the mean Shears weight/coefficient by patient and cell type with min 10 cells. UMAP plots indicate the position of cells positively (red) or negatively (blue) associated with the mutation. (**A**) Association of cellular composition with APC mutation in CRC patients. (**B**) Association of cellular composition with TP53 mutation in CRC patients.

**Figure S4:**
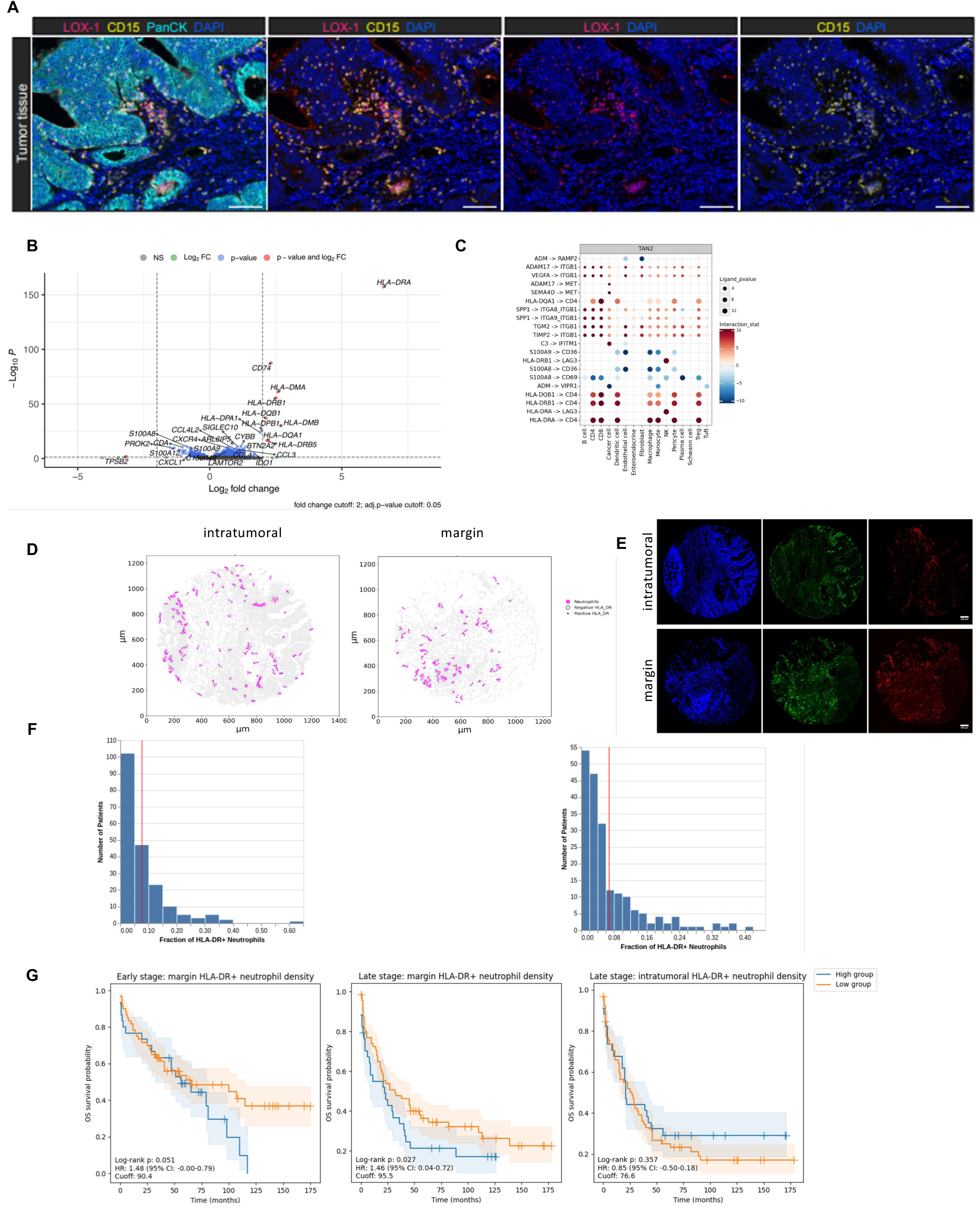
related to Figure 4. Characterization of neutrophils in CRC. (**A**) Selected multiplex immunofluorescence (mpIF) staining of LOX-1 (pink), CD15 (yellow), pan-cytokeratin (turquoise), and DAPI (blue) in colorectal cancer (CRC) tumor tissue. Scale bar: 100 µm. (**B**) Differentially expressed genes between HLA-DR^+^ and HLA-DR^-^ neutrophils paired per patient. (**C**) HLA-DR^+^ neutrophil interactions, with ligand-receptor pairs on the Y-axis and target cells on the X-axis. ligand_pvalue reflects ligand significance in DEA, while Interaction_stat integrates expression levels and statistical significance. (**D**) Neutrophil niches in the intratumoral and tumor margin regions detected by IF staining of CD15 in a CRC patient. (**E**) Images of IF stainings corresponding to (**D**), DAPI (blue), CD15 (green) and HLA-DR (red). (**F**) Fraction of HLA-DR^+^ neutrophils in tumors from CRC patients (left) and fraction of niche resident HLA-DR^+^ neutrophils in tumors from CRC patients (right). (**G**) Kaplan-Meier curves for overall survival stratified by high (upper 34%) and low HLA-DR^+^ neutrophil density (cells/mm^2^).

**Figure S5:**
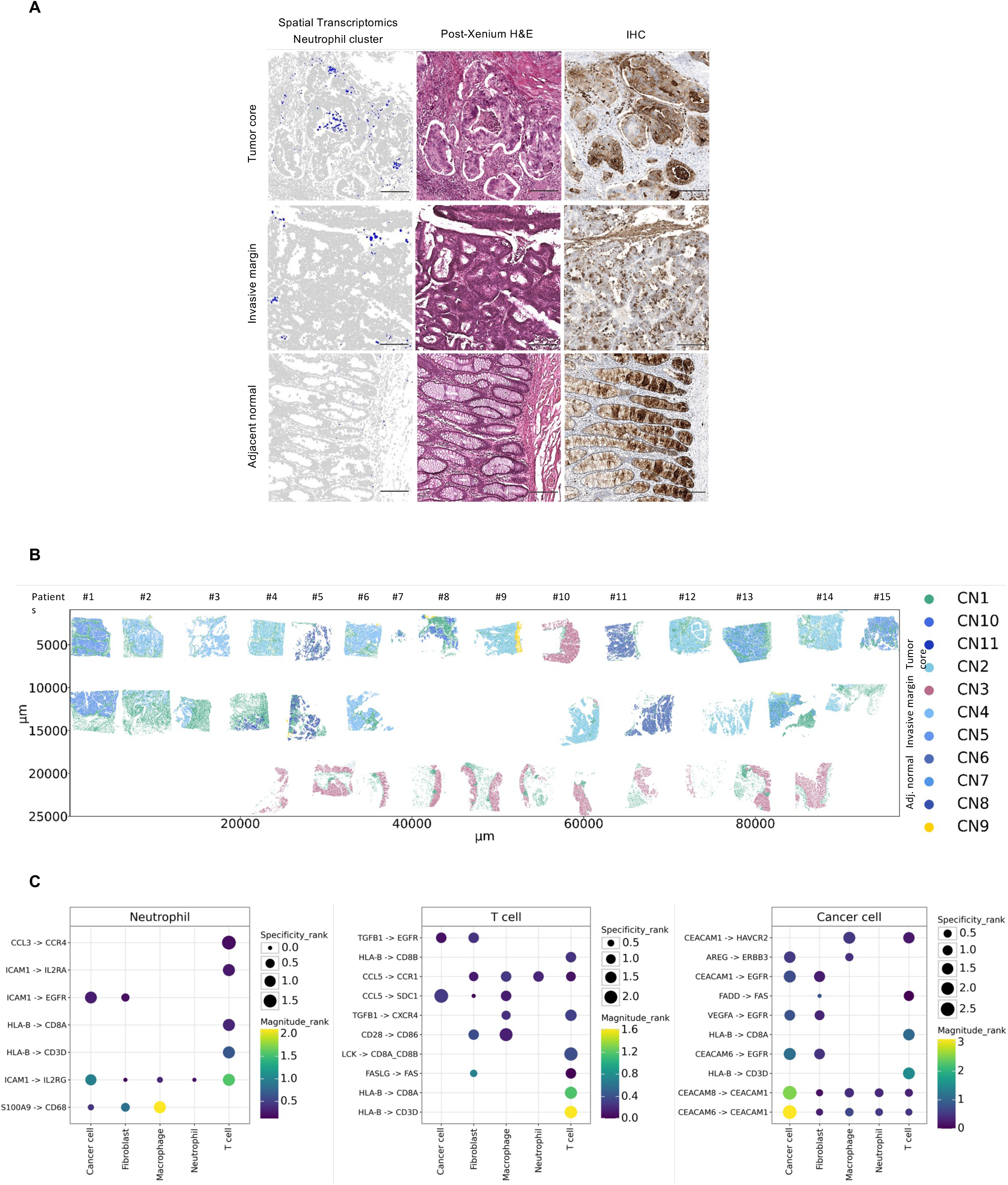
related to Figure 5. Spatial single-cell multi-modal profiling. (**A**) Representative images of neutrophil clusters in a CRC tumor core, invasive margin and adjacent normal tissue, annotated by spatial transcriptomics profiling (left panels). Matched H&E stainings (middle panels) and IHC stainings with CD15 AB (right panels) of consecutive FFPE slices were performed for validation and confirmed by pathologists. Scale bar: 200 µm. (**B**) Cellular niches defined in the spatial transcriptomics data (CN1: stromal niche; CN3: normal epithelial niche; CN9: neutrophil-dominated niche; CN2, CN4-8 and CN10-11 represent distinct cancerous niches). (**C**) Cell-cell-interaction pathways signalling from neutrophils, T cells and cancer cells, respectively, towards the main target cell populations in CN9 (specificity_rank: aggregated specificity score; magnitude_rank: aggregated magnitude score).

**Figure S6:**
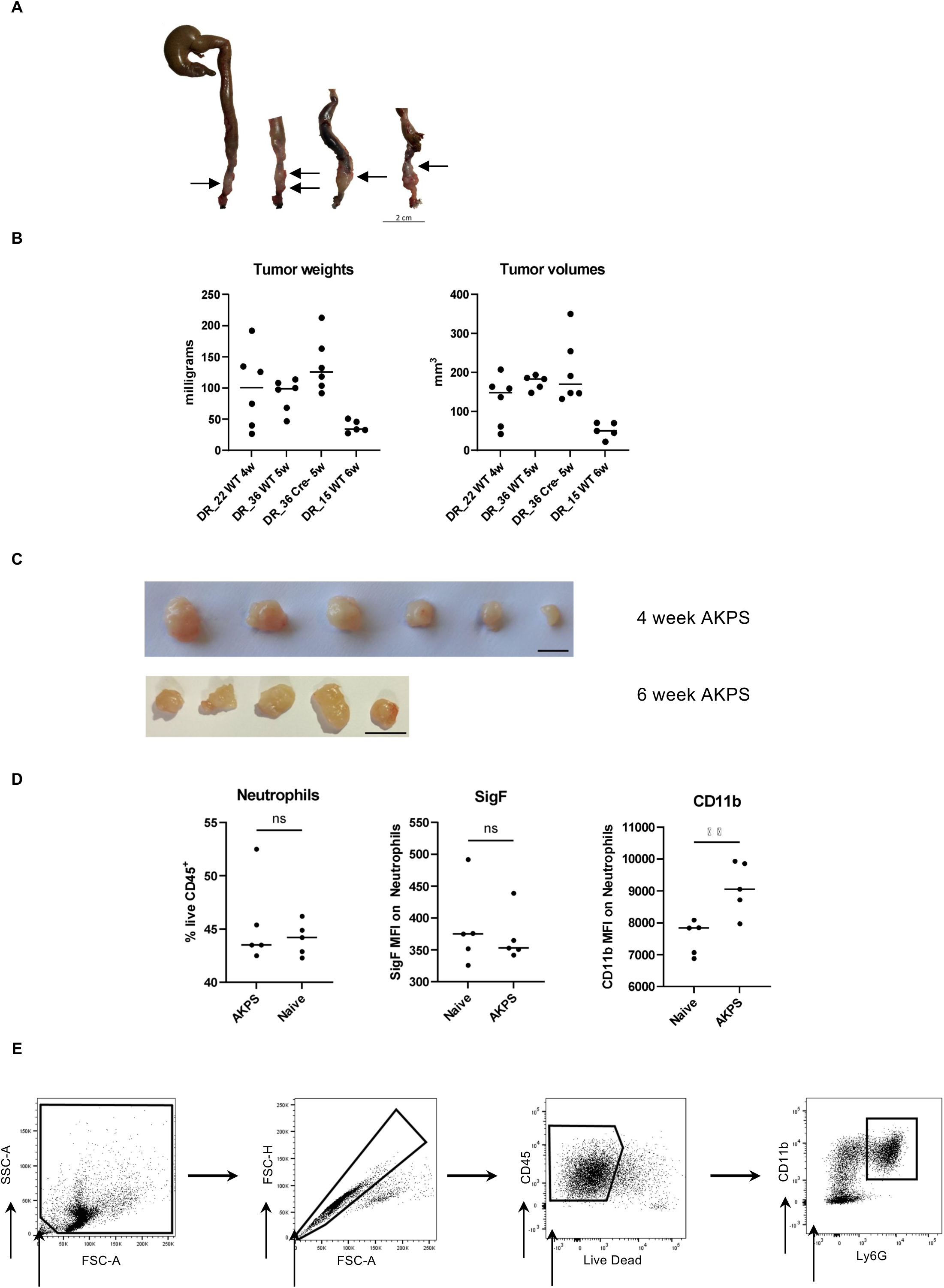
related to Figure 6. Neutrophils in a murine orthotopic organoid transplant model for CRC. (**A**) Orthotopic transplantation of AKPS organoids (location indicated through arrows). Scale bar: 2 cm. (**B**) Tumor weights and tumor volumes of the mice. (**C**) Picture of the respective tumors. Scale bar: 10 mm. (**D**) Neutrophils in the bone marrow of the mice after 6 weeks. (**E**) Representative plots of the flow cytometry gating strategy to define neutrophils in the bone marrow.

**Figure S7:**
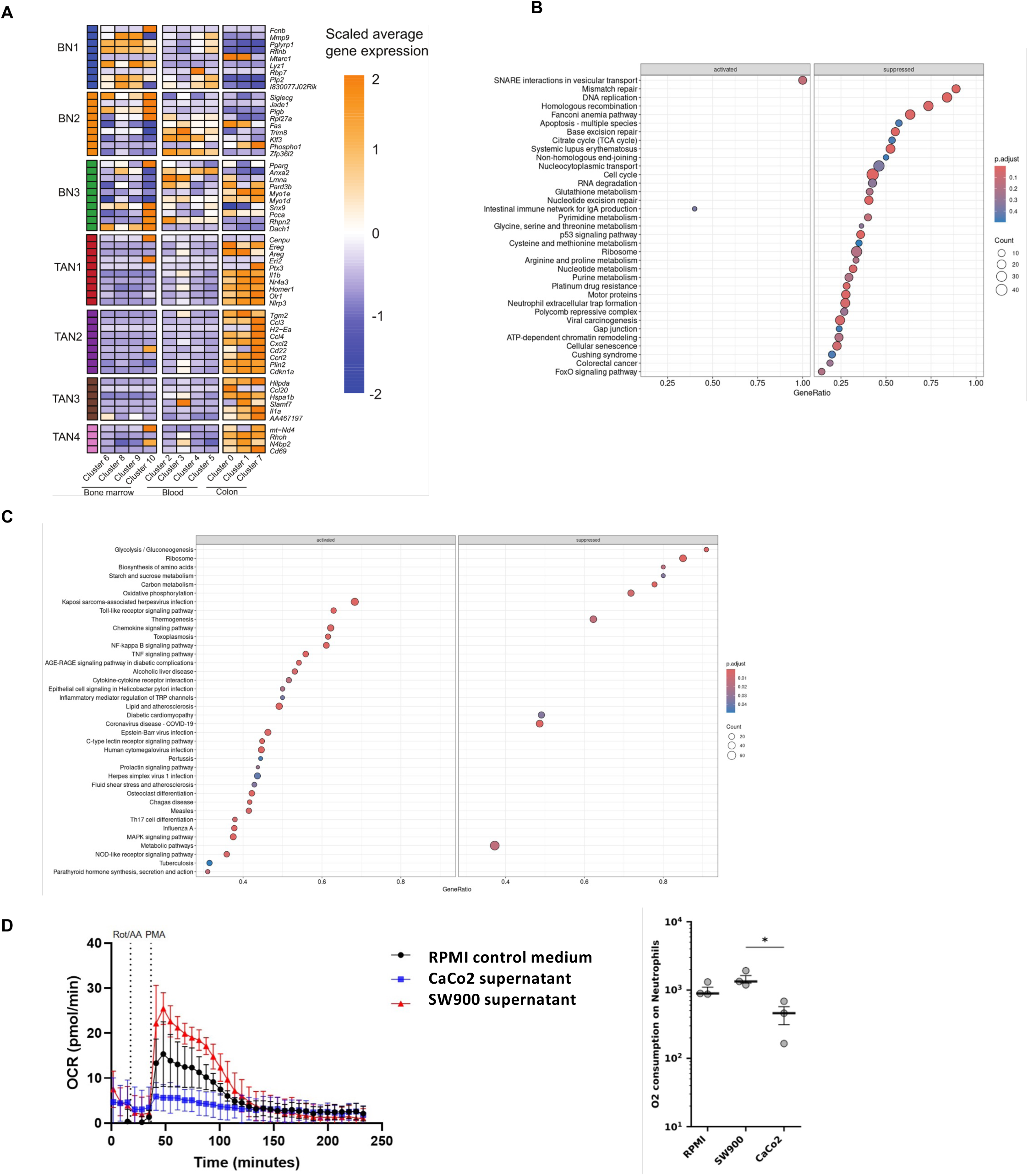
related to Figure 7. Neutrophil phenotypes along the bone marrow-tumor axis. (**A**) Heatmap showing the scaled average gene expression by cluster of the ortholog genes representing human neutrophil clusters. Columns are clustered by tissue presence. n = pool of 3 (non-tumor blood and bone marrow) or 5 (non-tumor colon and tissues from AKPS tumor-bearing mice) mice. Gene Set Enrichment Analysis (GSEA) on KEGG terms for (**B**) the expression profiles of bone marrow neutrophils from AKPC mice (n = pool of 5 mice) and controls (n = pool of 3 mice), and (**C**) the expression profiles of blood neutrophils from healthy volunteers and patients with CRC. (**D**) XF Neutrophil Activation Assay kinetic traces of oxygen consumption rate (OCR) of healthy human neutrophils after 24h culture in SW900 supernatant, CaCo2 supernatant or RPMI control medium. Mitochondrial function is blocked by Rot/AA injection, oxidative burst is activated by PMA injection. O_2_ consumption is calculated from area under the curve (AUC) of kinetic trace. The data shown are the mean and SD, n = 3 biological replicates.

**Figure S8:**
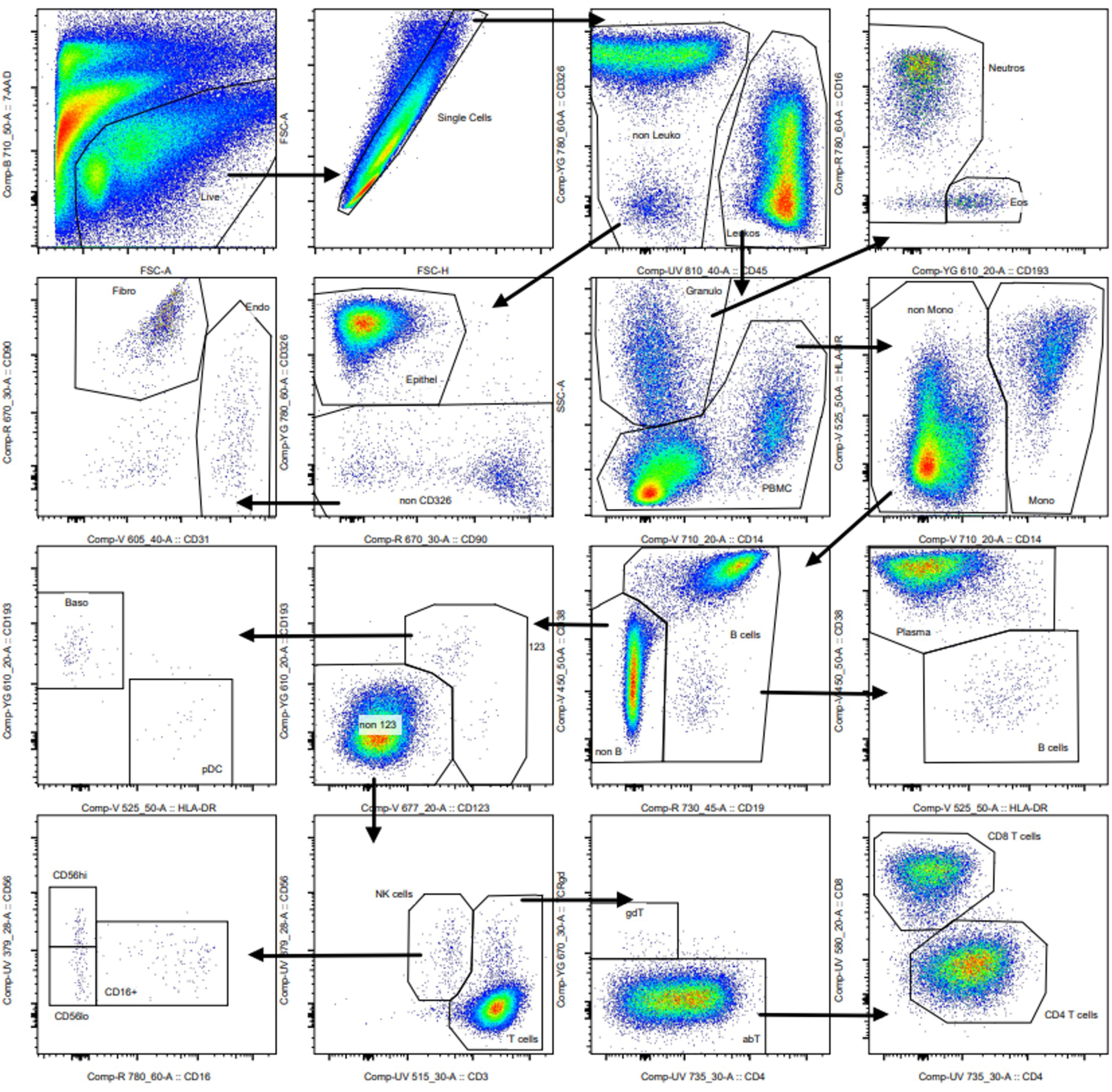
related to STAR Methods. Flow cytometry gating strategy to define cell populations from CRC tumor tissue, normal adjacent tissue and whole blood samples. In initial cleaning steps dead cells, debris and doublets were removed using 7-AAD staining and scatter characteristics. Leukocytes were defined by CD45 staining and sequentially gated into subtypes including neutrophils, monocytes, T cells and B cells. Non-CD45^+^ cells were gated into epithelial cells, endothelial cells and fibroblasts.

